# A Mechanism-based Outbreak Projection Study of Pertussis (Whooping Cough): Combining Particle Filtering and Compartmental Models with Pre-vaccination Surveillance data

**DOI:** 10.1101/598490

**Authors:** Xiaoyan Li, Nathaniel D. Osgood

## Abstract

Particle filtering is a contemporary Sequential Monte Carlo state inference and identification methodology that allows filtering of general non-Gaussian and non-linear models in light of time series of empirical observations. Several previous lines of research have demonstrated the capacity to effectively apply particle filtering to low-dimensional compartmental transmission models. We demonstrate here implementation and evaluation of particle filtering to more complex compartmental transmission models for pertussis – including application with models involving 1, 2, and 32 age groups and with two distinct functional forms for contact matrices – using over 35 years of monthly and annual pre-vaccination provincial data from the mid-western Canadian province. Following evaluation of the predictive accuracy of these four particle filtering models, we then performed prediction, intervention experiments and outbreak classification analysis based on the most accurate model. Using that model, we contribute the first full-paper description of particle filter-informed intervention evaluation in health. We conclude that applying particle filtering with relatively high-dimensional pertussis transmission models, and incorporating time series of reported counts, can serve as a valuable technique to assist public health authorities in predicting pertussis outbreak evolution and classify whether there will be an outbreak or not in the next month (Area under the ROC Curve of 0.9) in the context of even aggregate monthly incoming empirical data. Within this use, the particle filtering models can moreover perform counterfactual analysis of interventions to assist the public health authorities in intervention planning. With its grounding in an understanding of disease mechanisms and a representation of the latent state of the system, when compared with other emerging applications of artificial intelligence techniques in outbreak projection, this technique further offers the advantages of high explanatory value and support for investigation of counterfactual scenarios.

## 1. Introduction

Pertussis is a common childhood disease, which is a highly contagious disease of the respiratory tract that caused by the bacterium *Bordetella pertussis* [1]. It is most dangerous for infants, due to risks of severe complications, post-paroxysm apnia [1]. The most frequent complication is pneumonia, while seizures and encephalopathy occur more rarely [1]. Pertussis is a highly contagious disease only found in humans, and spreads from person to person by coughing, sneezing, and prolonged proximity [2]. Evidence indicates a secondary attack rate of 80% among susceptible household contacts [3]. In contrast to some other prevalent childhood diseases, immunity conferred by natural exposure or vaccination to pertussis is widely believed to wane relatively rapidly, leading to significant risks of infection even in adults who have been previously infected. It is notable that babies can be infected by adults, such as parents, older siblings, and caregivers who might not even know they have already contracted this disease [2]. Pertussis incidence shows no distinct seasonal pattern. However, it may increase in the summer and fall [3].

In the pre-vaccination era, pertussis was one of the most common childhood infectious diseases and a major cause of childhood mortality. In 1860, the mortality rate of all-age pertussis in Demark was 0.015% [4], but that burden fell heavily on infants and children. Research into historical mortality rates from pertussis indicate that the death rate in infancy is higher than in other groups [4]. In recent years globally, there are an estimated 24.1 million cases of pertussis, and about 160,700 deaths per year [5]. Since the 1980s, there has been a rising trend in the reported cases of pertussis in the United States [5]. The most recent peak year of the reported cases of pertussis in the United States is 2012, when the Centers for Disease Control and Prevention (CDC) reported 48,277 cases, but many more are believed to go undiagnosed and unreported [5]. Research aimed at estimating the level of population susceptibility and predicting the transmission dynamics of pertussis could aid outbreak prevention and control efforts by health agencies, such as performing intervention before the predicted next outbreak, and in targeted outbreak response immunization campaigns [6].

Dynamic modelling has long served as an important tool for understanding the spread of the infectious diseases in population [7], including pertussis, and for evaluating the impacts of interventions such as immunization and hygeine-enhancing. In recent years, particle filtering as a machine learning method has been employed for incorporating empirical time series data (such as surveillance [8] and online communicational behavior data [9]) to ground the hypothesis as to the underlying model state models in some previous researches [10, 11, 12, 13], especially for the infectious diseases of influenza [14, 15] and measles [8]. In this paper, we apply the particle filtering algorithm in a more complex and widely used compartmental model [16] of pertussis by incorporating the reported pertussis cases in Saskatchewan during the pre-vaccination era. Particle filtering for pertussis is different than for other pathogens on account of the need for state estimation to estimate the population segments at varying levels of immunity. Another need concerns extends from the heterogenous nature of the mixing and incidence burden between different age groups. For this reason, age-structured models are examined here. Specifically, we have examined two categories of age-structured particle filtering models – with 2 age groups and with 32 age groups. Moreover, we have proposed and explored three methods for calculating the contact matrix, so as to reduce the degrees of freedom associated with characterization of the contact matrix. This contribution compares the results obtained from all the particle filtering models by incorporating the empirical data across the whole timeframe evaluating the predictive accuracy of the models. Finally, using the minimum discrepancy particle filtering model, we demonstrate how we can evaluate intervention effects in a fashion that leverages the capacity of particle filtering to perform state estimation.

## 2. Methods and materials

### 2.1. Mathematical epidemiological models

As noted above, the dynamics of pertussis in the population is more complex than for infectious diseases that confer lifelong immunity – including other prominent childhood infectious diseases such as measles – due to the temporary character of the immunity acquired by *Bordetella pertussis* infection. As the time since the most recent pertussis infection increases, the immunity of a person wanes [7]. People with lower immunity generally tend to be more easily infected, and exhibit a higher risk of transmitting the infection once infected.

In this paper, we have employed the structure of the popular pertussis mathematical model of Hethcote [16]. To capture the characteristics of pertussis in waning of immunity and the different level of infectiousness and susceptibility involved with infection in light of pre-existing immunity, the compartmental model in [16] further divides the infectious population into three groups: infective with weak-disease (*I*_*w*_), mild-disease (*I*_*m*_), and full-disease (*I*). In a similar fashion, the recovered population is divided into four groups of successively increasing immune system strength: *R*_1_, *R*_2_, *R*_3_ and *R*_4_.

Figure 1 shows the mathematical structure of our compartmental pertussis model adapted from [16]; readers interested in further introduction of this structure are referred to Appendix A. It is notable that the model of Hethcote (1997) [16] employs a formulation in which each state variable is of unit dimension, representing a fraction of the population in different age groups of the same class. However, for the sake of easing comparison against empirical data – the pertussis reported cases in the province of Saskatchewan, Canada during pre-vaccination era (from 1921 to 1956) – two parts are modified compared to the original model in [16]. Firstly, the model in this paper is represented in a re-dimensionalized fashion, with the state variables representing counts of persons based on the structure in Figure 1. Secondly, because of the focus of this paper on the pre-vaccination error, all vaccinated-related elements of the original model of [16] are removed.

**Figure 1:**
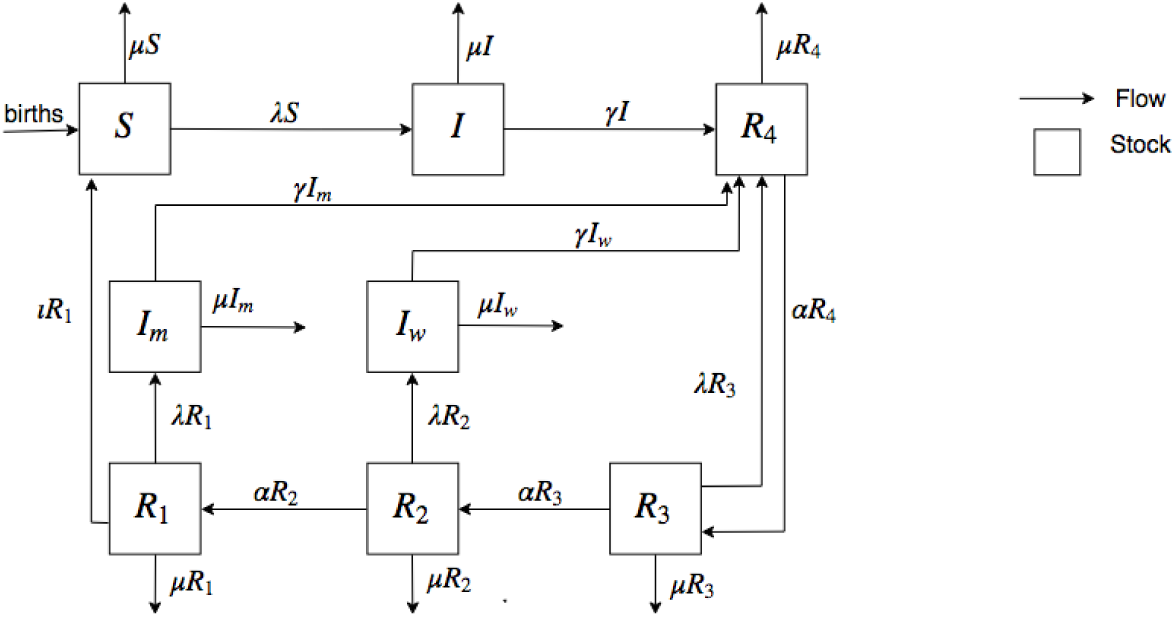
The transfer diagram for the pertussis model without vaccination. adapted from [16]

Finally, four models are considered in this research. Using *n* to denote the count of age groups incorporated in the models, we consider models of the aggregate population (*n* = 1), of two age groups (*n* = 2), 32 age groups model (*n* = 32) with the contact matrix introduced in the paper of Hethcote (1997) [16], and a final model with 32 age groups (*n* = 32) model with a re-balanced contact matrix. The mathematical models are introduced in separate sections below.

#### 2.1.1. *Aggregate population epidemiological model (n* = 1)

In the aggregate model, discordant contacts – contacts of infectious individuals (including the persons in stocks of *I, I*_*m*_ and *I*_*w*_) and the others (including the persons in the other stocks, *S, R*_1_, *R*_2_, *R*_3_ and *R*_4_) – are mixed homogeneously. Based on the mathematical structure (Figure 1) adapted from Hethcote (1997) [16], the equations of the aggregate compartmental model of pertussis are as follows:

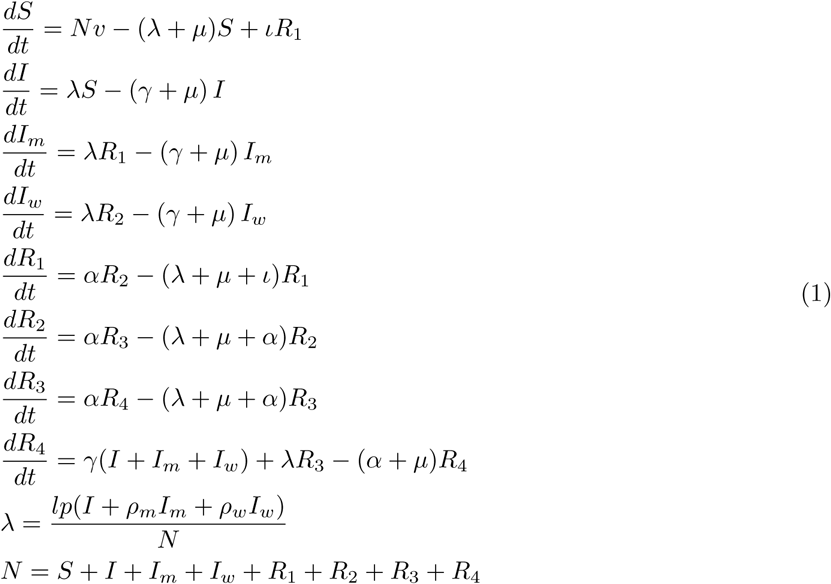

The meaning of the states and parameters are as follows: Compartment *S* is the count of susceptible individuals. Compartments *I, I*_*m*_ and *I*_*w*_ are the count of individuals having full-disease infectious cases with all of the usual symptoms, with mild disease and weak disease infectious cases, respectively, and with correspondingly decreasing infectivity. It is notable that individuals in both class *I*_*m*_ and *I*_*w*_ lack usual symptoms of pertussis, and thus exhibit atypical pertussis [16]. Compartments *R*_1_, *R*_2_, *R*_3_ and *R*_4_ are the count of recovered people in the population, with correspondingly increasing levels of immunity. *N* is the population size. *v* is the overall population birth rate, while *µ* is the death rate. It is notable that this paper follows [16] in assuming that all model compartments share identical values for the mortality rate (*µ*), although the death rates in the stocks of the infectives (*I*_*m*_, *I*_*w*_, and especially *I*) are theoretically higher than for the other stocks, due to risk of pertussis-induced mortality. The mean time for waning of immunity from the stock of *R*_1_ to *S*, and that for successive waning of immunity from successive pairs of *R*_3_, *R*_2_, and *R*_1_, are *ι*^−1^ and *α*^−1^, respectively. The three infectious compartments – *I, I*_*m*_, *I*_*w*_ – share an identical mean infectious periods of *γ*^−1^. However, the infectiousness of an individual varies across the three infectious compartments (*I, I*_*m*_, *I*_*w*_), with individuals in compartment *I, I*_*m*_ and *I*_*w*_ having highest, middle and lowest infectiousness, respectively. Parameters *ρ*_*m*_ and *ρ*_*w*_ represent the ratios of the infectiousness of those in the mild-disease (*I*_*m*_) and weak-disease (*I*_*w*_) infectious classes to those in the full-disease infectious classes of *I* [16]. The force of infection parameter *λ* characterizes the hazard rate – the probability density with which a susceptible (a person in the stocks of *S, R*_1_, *R*_2_ and *R*_3_) is subject to infection from an infective, and is governed by the mass action principles [17, 16]. Parameter *λ* is related to the total effectively infectious cases (*I* + *ρ*_*m*_*I*_*m*_ + *ρ*_*w*_*I*_*w*_), contact rate (denoted as *l*) and per-discordant-contact transmission probability (denoted as *p*).

#### 2.1.2. General age-structured epidemiological model

To capture the difference of the contact pattern among different age groups – for example, the fact that children in school age primarily contact with peers, while babies contact more closely with their parents or caregivers – and adapt the simulation models with the empirical datasets (both monthly pertussis reported cases across the whole population and age-group-specific yearly pertussis reported cases), we extended the pertussis model in Equation (1) to an age-structured model.

##### The age-structured demographic model

Before introducing the epidemiological age-structured pertussis mathematical model, we first introduce the age-structured demographic model. The demographic model mainly captures the age structure and the birth and death in the population related to the empirical data (pertussis reported cases in province of Saskatchewan in Canada during the pre-vaccination era – from 1921 to 1956) employed in this paper. Suppose we have *n* age groups in the whole population, by divided by a sequence of ages *a*_*i*_, 1 ≤ *i* ≤ *n* − 1. The age groups can be characterized as a series of *n* intervals – [0, *a*_1_), [*a*_1_, *a*_2_), …, [*a*_*n*−1_, ∞). The demographic model can then be written as follows [8, 16, 18].

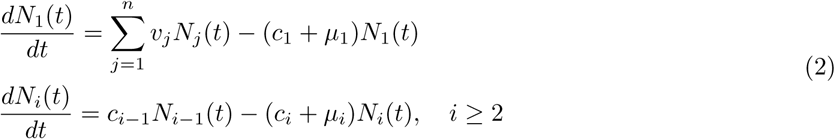

where *N*_*i*_ is the number of people in age group *i*; *v*_*i*_ and *µ*_*i*_ are the birth and death rate of age group *i*, respectively; *c*_*i*_ is the aging rate of age group *i*, given by *c*_*i*_ = (*a*_*i*_ − *a*_*i*−1_)^−1^, and *c*_*n*_ = 0.

In this paper, we assume that the population is in equilibrium; this reflects the fact that the empirical Saskatchewan population size from 1921 to 1956 does not change dramatically [19], as will be discussed below in greater detail. This approximation assumes that the total population *N*_*i*_(*t*) of age group *i* will remain invariant over the model time horizon, that is, *dN*_*i*_(*t*)*/dt* = 0. Thus, according to Equation (2), for this simplified context, the death rate *µ*_*i*_ can be calculated as follows:

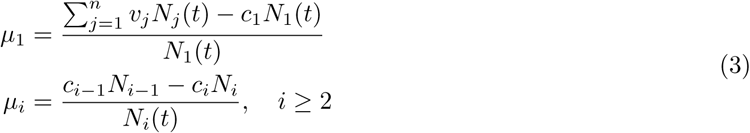

The values of parameters in the demographic model are estimated from the empirical data. Specifically, the population in each age group *N*_*i*_ is estimated from the age pyramid of Saskatchewan [19], and the birth rates *v*_*i*_ are estimated from the Public Health Annual Report of Saskatchewan [20] published yearly by the Government of Saskatchewan.

##### The age-structured pertussis epidemiological model

By incorporating the age-structured demographic model shown in Equation (2), and the structure of the compartmental epidemiological model shown in Figure 1, we obtain the age-structured pertussis epidemiological model given below. Readers interested in the detailed mathematical derivation are referred to our previous contribution [8].

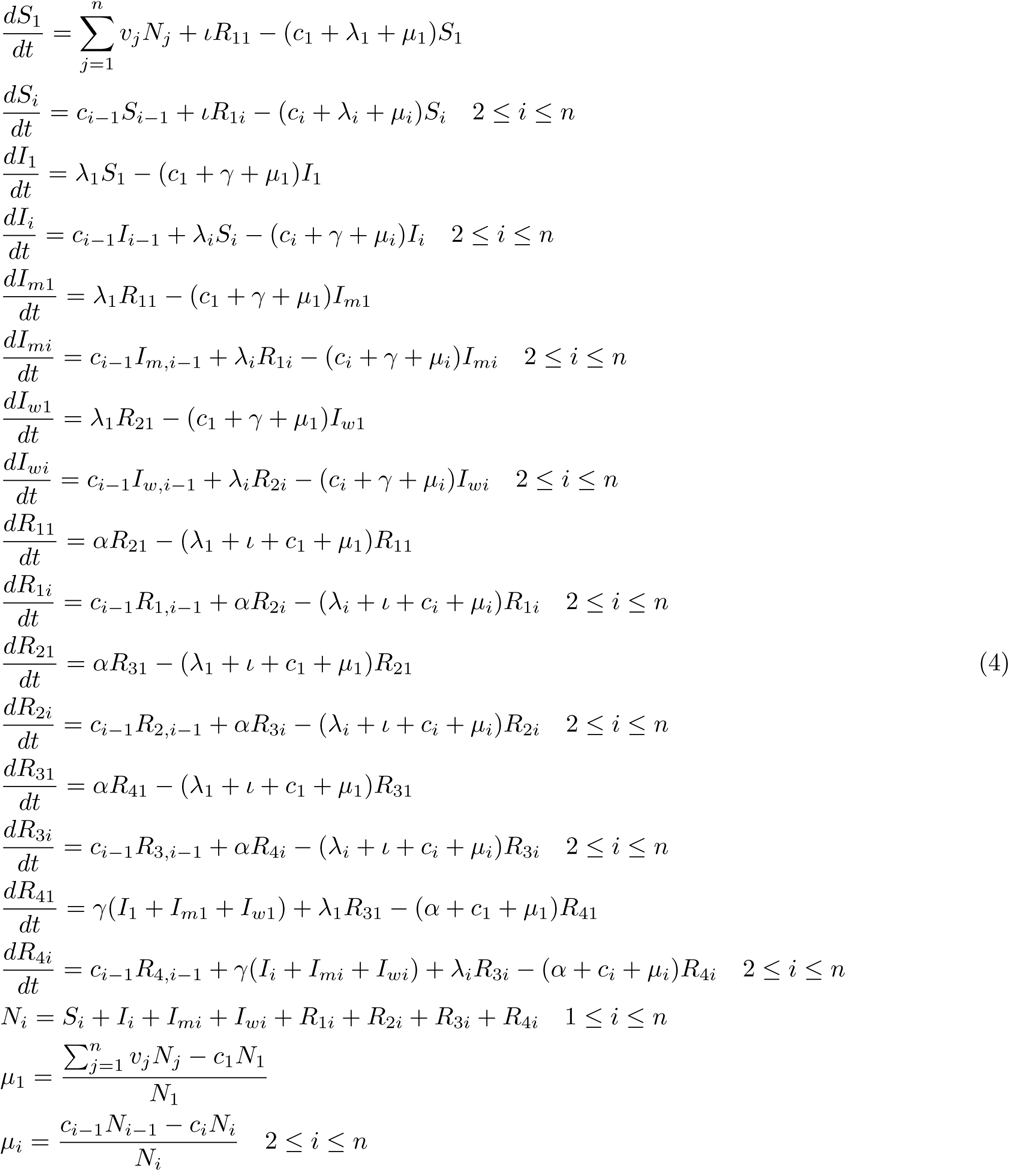

In this age-structured epidemiological model, the definition of most quantities are consistent with (*mutatis mutandis*) the aggregate population epidemiological model (Equation (1)) and the age-structured demographic model (Equation (3)), with the notable exception of the force of infection *λ*_*i*_ for age group *i*.

As noted above, this work followed [16] in characterizing transmission of pertussis infection between infectives and susceptibles according to mass action principles. The force of infection defined as the hazard rate with which susceptibles are infected by infectives, and is related to contact rate, transmission probability, and the fraction of infectives in the whole population. In the model with aggregate population, the individuals are assumed contact homogeneously, and the force of infection can be simply calculated as in Equation (1). However, in the age-structured model, contacts between individuals are assumed to occur homogeneously within age groups and heterogeneously across different age groups. Thus, the calculation of force of infection in the age-structured models are considerably more complex than for aggregate population model, being mediated by a contact matrix. Readers interested in the mathematical representation of the contact matrix could are referred to our previous contribution [8]. In the current paper, we have employed three different methods of calculating the contact matrix and – by extension – the force of infection in the models. These three methods are introduced as follows.

#### 2.1.3. Force of infection models

##### General mass action-based contact matrix

Under the assumption of mass action, the force of infection – the hazard rate (probability density) with which a susceptible is transmitted the pathogen by infectives – can be calculated by the sum of the hazard rates associated with transmission from infectives in each age group in turn. The force of infection of each age group is correspondingly represented as follows.

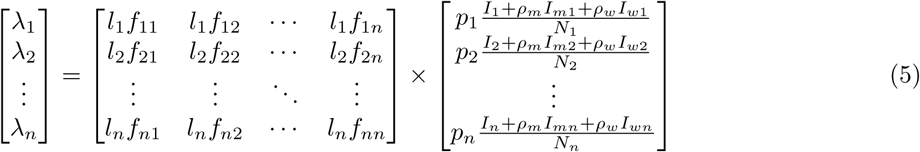

The above can be rewritten as the following equation:

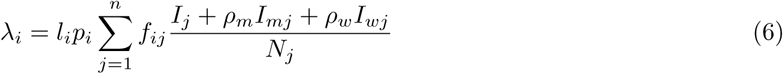

where *λ*_*i*_, *l*_*i*_, *p*_*i*_, *I*_*i*_, *I*_*mi*_ and *I*_*wi*_ are the force of infection, contact rate, transmission probability, number of persons in full-disease infectious, number of persons in mild-disease and weak-disease infectious classes in age group *i*, respectively. For an individual in age group *i, f*_*ij*_ is the fraction of that individual’s contacts that occur with others in the age group of *j*. Thus, for a given age group *i*: 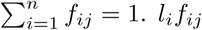 are then the elements in the contact matrix.

An advantage of this method in calculating the contact matrix in the age-structured model is that the contacts between any two age groups (e.g., *i* and *j*) is balanced (symmetric) – the number of total contacts of an age group *i* to group *j* equals to the number of total contacts of the age group *j* to group *i*; that is, *N*_*i*_*l*_*i*_*f*_*ij*_ = *N*_*j*_*l*_*j*_*f*_*ji*_. However, this method has a notable disadvantage that the count of unknown parameters in calculating the contact matrix grows quadratically with the count of age groups (denoted as *n*) in the model; a demonstration of the super-linear growth of the total number of unknown parameters in the contact matrix with the total number of age groups is shown in Appendix B. This disadvantage makes challenging parameter estimation for models incorporating a large number of age groups. To address this challenge, we have explored two other methods for characterizing the contact matrix and force of infection in which the count of parameters grows sub-linearly or linearly with the total number of age groups. The first is a method of obtaining an un-balanced contact matrix contributed by Hethcote (1997) with a constant number of unknown parameters [16]. The second approach calculates a re-balanced contact matrix in which the number of unknown parameters grows linearly with the total number of age groups. Each of these approaches are characterized below.

##### The Unbalanced Contact Matrix

This unbalanced contact matrix is introduced in the research of Hethcote (1997) [16], which assumes that only adequate contacts are sufficient to transmit the disease. This method based on a simple proportional mixing assumption that the number of total persons contacted by one person in the age group *j* is distributed among the population in the age group *i* in proportion to the fractions *l*_*i*_*/D*\*, where *D*\* is the total number of contacts per unit time received by all people, *l*_*i*_ is the contact rate – average number of persons contacted by a person per unit time – of age group *i*, and 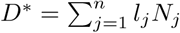 [16]. The elements in the contact matrix are *l*_*i*_*l*_*j*_*/D*\* [16]. Finally, the re-dimensionalized force of infection (*λ*) used in Equation (4) and in [16] is given as follows:

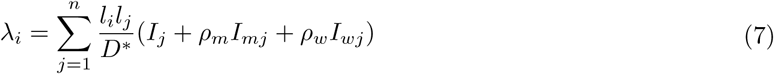

However, in this paper, we employ the dimensionless representation of the “force of infection” in Equation (8), which is consistent with [16], instead of the re-dimensionalized one in Equation (7). The motivation for this lies in our use of the values of parameters related to the mixing matrix from [16], which will be detailed below in the section “particle filtering implementation”. The dimensionless equation of force of infection in [16] is as follows:

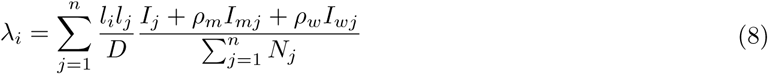

where *D* is the dimensionaless total contacts across all population, and 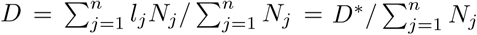.

The advantage of this method is that – if one adopts the values of the contact rate in each age group given in [16] – there are no unknown parameters required for calculating the contact matrix. And it is straightforward to calculate the contact matrix as long as those age-specific contact rate parameters are known. However, this method of calculating the contact matrix suffers from a notable disadvantage – a lack of guaranteed symmetry between the contacts exerted between pairs of age groups. Specifically, it can be readily shown that the value of the total contacts occurring from age group *i* to age group *j* is not in general equal to the value of the total contacts occurring from age group *j* to age group *i*. This reflects the fact that the number of total contacts of the age group *j* to age group *k* is *N*_*j*_*l*_*j*_*l*_*k*_*/D*\*, while the number of total contacts of the age group *k* to age group *j* is *N*_*k*_*l*_*j*_*l*_*k*_*/D*\*. In general, these two quantities need not be equal.

To address this shortcoming, we explored a previously contributed method to calculate a balanced contact matrix. While the above method does not require additional parameters, for the balanced method, the total number of the unknown parameters grows linearly with the number of age groups.

##### The Re-balanced Contact Matrix

To calculate the balanced contact matrix, we have employed the method introduced in research by Garnett and Bowden (2000) [21]. The elements of the contact matrix *l*_*ij*_*f*_*ij*_ and force of infection *λ*_*i*_ are as follows; readers interested in the detailed mathematical deduction of the re-balanced contact matrix can refer to Appendix C:

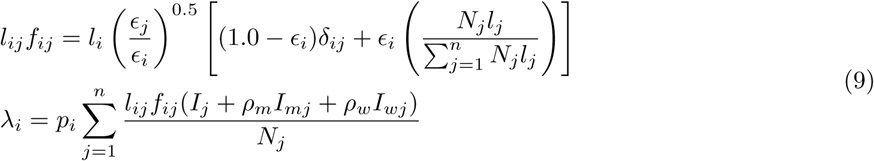

where *p*_*i*_ is the transmission probability of age group *i, f*_*ij*_ is the fraction of the contacts of an individual in age group *i* that are made with others in age group *j, δ*_*ij*_ is the identity matrix, mixing parameter *ϵ*_*i*_ determines where mixing occurs on a scale from fully homophilic – persons only contact with the individuals in the same age group (representing *ϵ*_*i*_ = 0) – to random mixing in which the contact among the total population is non-preferential (representing *ϵ*_*i*_ = 1.0).

Finally, based on the above discussion, we have employed four pertussis epidemiological models as the state-space models to be used in corresponding applications of particle filtering – the aggregate population model (shown in Equation (1)), the age-structured model with two age groups (Equation (4) with *n* = 2) with the general contact matrix based the mass action assumption (Equation (6)), the age-structured model with 32 age groups (Equation (4) with *n* = 32) with an un-balanced contact matrix (Equation (8)) and the age-structured model with 32 age groups (Equation (4) with *n* = 32) with a re-balanced contact matrix (Equation (9)). It is notable that the 32 age group division applied is directly adopted from Hethcote (1997) [16], with age groups from 0–1 month, 2–3 months, 4–5 months, 6–11 months, integer ages for 1 through and including 19, 20–24 years, 25–29 years, 30–39 years, 40–49 years, 50–59 years, 60–69 years, 70–79 years, 80–89 years, 90 years and older. The age structured model with two age groups dichotomizes the population into 0–4 years and 5 years and older age categories. It bears noting that while more detailed age structure can better capture both the effects of population aging and inter-group heterogeneity, in terms of particle filtering, it entails estimation of a larger underlying model state space – potentially adversely affecting the accuracy of that estimation; in many models, it also requires specification of additional parameter values.

### 2.2. Particle Filter Implementation

Particle filtering is a contemporary state inference and identification methodology that allows filtering of general non-Gaussian and non-linear state space models in light of time series of empirical observations [10, 11, 22, 15, 8, 23]. This approach estimates time-evolving internal states of dynamic systems where random perturbations are present, and information about the state is obtained via noisy measurements made at each time step. The state space model characterizes the processes governing evolution over time of internal states with stochastics consisting of random perturbations. The states in the state space model are assumed in general to be latent and unobservable. Information concerning the latent states is obtained from a noisy observation vector. The means by which the particle filter method operates includes the Recursive Bayes Filter [22], Sequential Importance Sampling [22, 23, 10], and Resampling [22, 23, 10].

Sequential importance sampling (SIS) is the most basic Monte Carlo method used to sample when the predict-and-update equations of the recursive Bayes filter are not analytically tractable [22]. The key idea of SIS is to estimate the posterior distribution at a given time with a weighted set of samples. SIS then recursively updates these prior samples to obtain samples approximating the posterior distribution at the next time step. These importance-weighted samples are also named particles [22]. The SIS particle filter commonly suffers from a strong degeneracy problem – as the algorithm continues, many – and eventually most – particles will develop a negligible weight. This occurs because we are sampling in a high dimensional space, using a myopic proposal distribution [23].

The key idea underlying resampling is a variant of the principle of “survival of the fittest”. To achieve this, the resampling step will monitor the effective sample size following each observational update. Whenever the effective sample size drops below a threshold, the algorithm will draw a new set of particles from the existing set, where the probability of drawing a given particle is – in accordance with the principle of importance sampling – proportional to its weight. Within such resampling, particles with higher weight will tend to be reproduced, and particles with lower weight will tend to die out. The new particles inherit their parent’s values but carry a uniform normalized weight. At a given time, each particle contributing to the distribution (represented collectively by the particles according to the principles of sequential importance sampling [23]) can be seen as representing a competing hypothesis concerning the underlying state of the system at that time. The particle filtering method can be viewed as undertaking a “survival of the fittest” of these hypotheses, with fitness of a given particle being determined by the consistency between the expectations of the hypothesis associated with that particle and the empirical observations.

Interested readers are referred to more detailed treatment in [22, 24, 23, 10].

#### 2.2.1. State Space Model

The state space model depicts the processes governing the state – both latent and observable – of a noisy system evolving with time. In this paper, we employ the deterministic pertussis epidemiological models as base models. Reflecting the fact that particle filtering offers value in the context of underlying state equation models exhibiting stochastic variability, we then extend these deterministic models by adding random perturbations in some processes or parameters, so as to represent the stochastic processes in the real world; the extended, stochastic model then serves as the basis for a corresponding particle filter. Thus, we have built four particle filtering models based on the respective pertussis compartmental epidemiological models introduced previously – the aggregate population model, two-age group model with the general contact matrix, and the 32-age group models with both un-balanced contact matrix and re-balanced contact matrix.

##### Stochastic Adaptation of the Aggregate Population State Space Model

In the aggregate population state space model, we employ the aggregate population pertussis compartmental epidemiological model (equation (1)) as the base model. Stochastics are added to this base model in three areas – in the rate of new infections, for the contact process between susceptibles and infectives, and in the reporting process for infected cases. The mathematical structure of the pertussis aggregate population state space model is shown in Figure 2. The stochastics associated with these factors represents a composite of two factors. Firstly, there is expected to both be stochastic variability in the pertussis infection processes and some evolution in the underlying transmission dynamics in terms of an evolving reporting rate, as well as changes in mixing. Secondly, such stochastic variability allows characterization of uncertainty associated with respect to model dynamics—reflecting the fact that both the observations and the model dynamics share a high degree of fallability. Given an otherwise deterministic simulation model such as that considered here, there is a particular need to incorporate added stochastic variability in parameters and flows to provide the model with the requisite openness to correction when observing a new empirical datum [8].

**Figure 2:**
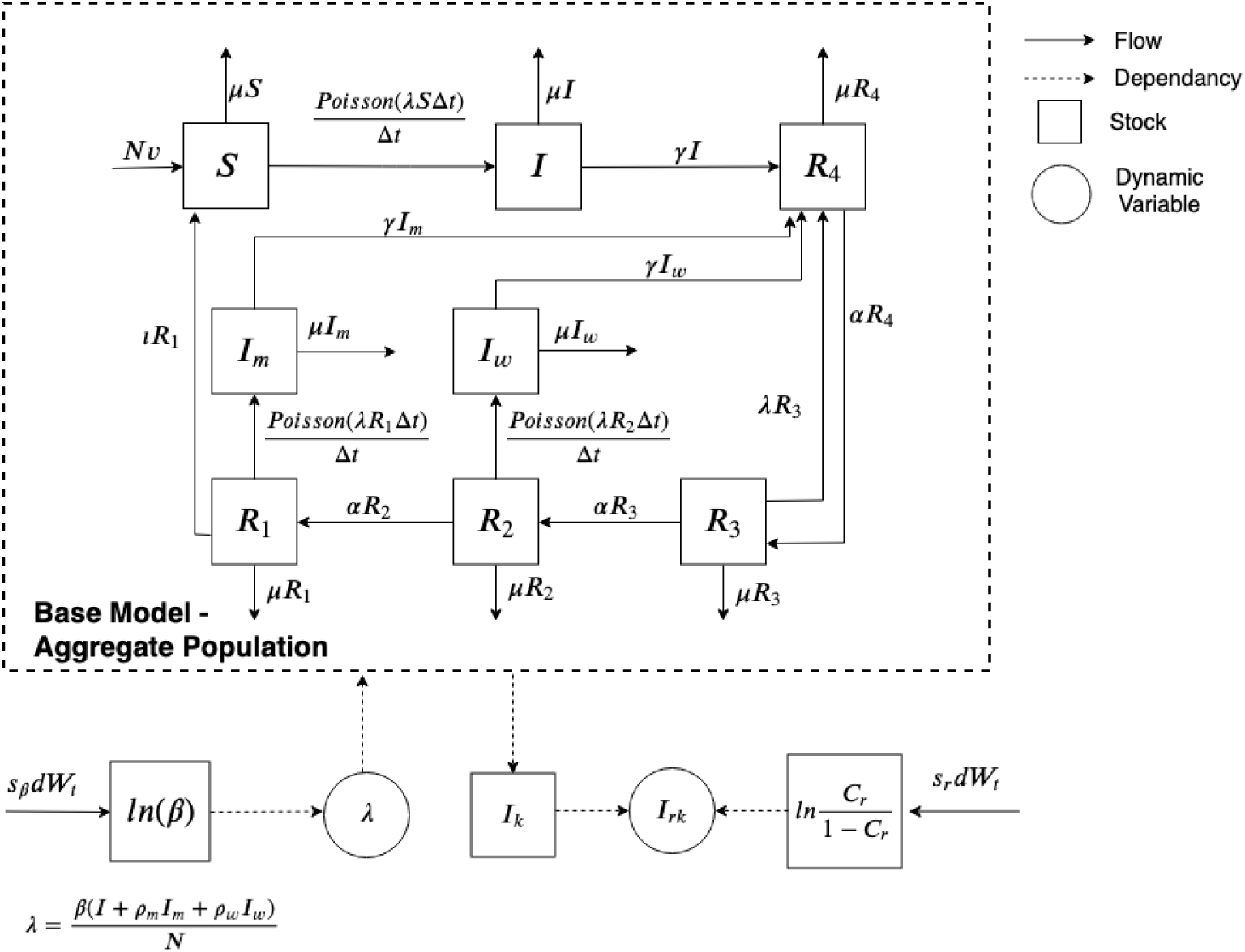
The mathematical structure of the aggregate particle filtered model.

In characterizing transmission process, we consider a stochastic process – specifically, a Poisson process – associated with incidence of infection, including cases of full-disease infectives (*I*), mild-disease infectives (*I*_*m*_) and weak-disease infectives (*I*_*w*_). This process reflects the small number of cases that occur over each small unit of time – denoted as Δ*t* (carrying the value of 0.01 months in all models in this paper, or roughly 7.3 days) [10, 8]. The new infection flows incorporating stochastic process (Poisson process) are correspondingly listed as follows:

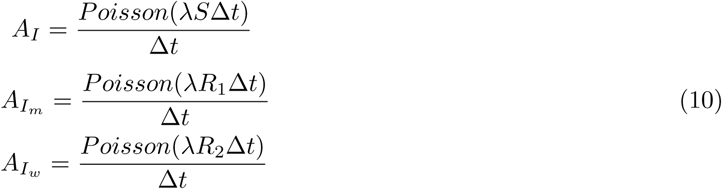

The second stochastic process considered in the aggregate population pertussis state space model is the mixing process between susceptibles and the infectious. We know that the transmission probability of the disease of pertussis is normally a constant. Thus, to simplify the model, we incorporated an effective contact rate parameter, denoted as *β*, where the effective contact rate is the multiplication of a per-month contact rate and transmission probability (of unit dimension), denoted as *l* and *p* (*β* = *lp*) in the deterministic aggregate population compartmental pertussis epidemiological model characterized in Equations (1). We posit that parameter will undergo some evolution in value in accordance with contact rates – such as due to social distancing, as the school year starts or stops, and enhanced hygenic awareness during outbreaks. We thus characterized effective contact rate *β* as evolving stochastically within the model.

To estimate changing values of the stochastic effective contact rate parameter *β*, and to investigate the capacity of the particle filter to adapt to parameters whose effective values evolve over simulation, we incorporated the parameter *β* into the state of the particle filter model, as seen in Figure 2. Moreover, reflecting the fact that the effective contact rate *β* is conceptually bounded to the non-negative real numbers, we treat the natural logarithm of the effective contact rate *β* as undergoing a random walk according to Brownian Motion, as characterized by a Wiener Process [25, 26, 8]. The stochastic differential equation of the effective contact rate *β* can thus be described according to Stratonovich notation as:

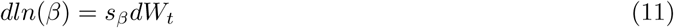

where *dW*_*t*_ is a standard Wiener process whose perturbations follow a normal distribution with 0 of mean and unit rate of variance; *s*_*β*_ is the diffusion coefficient. Thus, the perturbations in the value of *ln*(*β*) are normally distributed with 0 of mean and variance *s*_*β*_ ^2^.

The third stochastic process considered in the noisy state space model relates to the reporting process for infected pertussis cases. Over the multi-decadal model time horizon (as circumscribed by the span of the empirical data from 1921 to 1956), and particularly on account of shifting risk perception, there can be notable evolution in the degree to which infected individuals or their guardians seek care. To capture this evolution, we incorporated another stochastically evolving parameter – the fraction of underlying pertussis cases that are reported (denoted as *C*_*r*_); as for the above parameters, this parameter is also treated as an element of evolving model state. Reflective of the fact that the reporting rate *C*_*r*_ is a probability limited to the range [0, 1], we characterize the logit of *C*_*r*_ as also undergoing Brownian Motion according to Stratonovich notation [8] as follows:

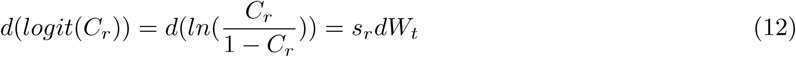

where *dW*_*t*_ is as above; *s*_*r*_ is the diffusion coefficient. Perturbations in the value of 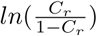 with time follow a normal distribution with mean 0 and variance *s*_*r*_^2^.

Moreover, to calculate the reported number of pertussis cases in the particle filtering model, which is used in the measurement model discussed below, we incorporated an extra state, denoted as *I*_*k*_, which accumulates the count of pertussis infectious cases from time *k* − 1 to time *k*. It is notable that the state of cumulative infectious cases from time *k* − 1 to *k* – *I*_*k*_ – is different from the original infectious states *I, I*_*m*_ or *I*_*w*_ in the deterministic compartmental model in Equations (1). Specifically, the state of the cumulative count of infectious cases *I*_*k*_ purely integrates all the inflows to the infectious states as a whole (and without all the outflows), so as to simulate a similar process of successively tallying up the pertussis cases over the course of some period of time as is undertaken in the real world. Moreover, we further assume that the individuals with mild-disease infectious cases (*I*_*m*_) and weak-disease infectious cases (*I*_*w*_) are also subject to reporting. The reporting rates of the mild-disease infectious cases (*I*_*m*_) and weak-disease infectious cases (*I*_*w*_) that have symptoms are considered to be *ρ*_*m*_ and *ρ*_*w*_, in this paper. It is notable that the sequence of the values of *k* correspond to the sequence of historical reporting times (per Month in this paper). Then, the cumulative infectious cases from time *k* − 1 to *k* in state *I*_*k*_ is represented as follows:

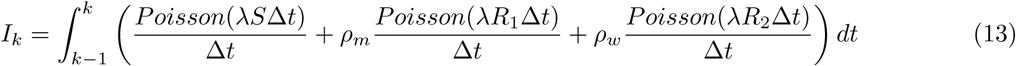

It bears emphasis that the model implementation of Equation (13) made use of identical values drawn for the stochastic components used in the flows that it serves to accumulate. Thus, the reported pertussis cases calculated in the state space model at the measure time *k*, denoted as *I*_*rk*_, can be represented as follows:

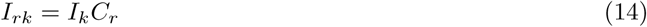

Finally, we obtained the noisy state space model of the pertussis particle-filtered aggregate model by incorporating into the base model – as given by the deterministic compartmental epidemiological model in equations of (1) – the adjusted stochastic parts in equations of (10), (11), (12), (13) and (14) (Figure 2). Readers interested in the complete mathematical equations, parameter values, and initial state assumptions for the state space model can refer to Appendix D; those seeking better understanding of basis for the parameters related to the transmission of pertussis in this model are referred to the research of Hethcote (1997) [16]. The demographic parameters of this model are sourced from the Annual Report of the Saskatchewan Department of Public Health [20] and the age pyramid of Saskatchewan [19]. The initial values of states in this model are estimated by tuning the particle-filtered model (the assumptions regarding the distribution of the initial states, as given by constants) and sampled by the particle filtering algorithm. Both the values of parameters and initial values of states in this model are listed in Appendix D.

##### The two-age group population structure state space model

In the two-age-group population structure state space model, we employ the age- and population-structured pertussis compartmental epidemiological model (Equation (4)) with *n* = 2 as the base model, where the variable of “force of infection” is calculated according to the mass-action based formulation of the general contact matrix (Equation (5)). In this model variant, we use subscripts “c” and “a” to denote the child- and adult-specific values, respectively, where the child age group includes all individuals from newborns to the end of the fourth year, and the remaining individuals are in the adult age group. Similar to the state space model with an aggregate population, noise is imparted to this base model in three elements – the new infectious occurrence process, the contact process between susceptibles and infectives, and the reporting process for infected cases. The mathematical structure of the pertussis aggregate population state space model is shown in Figure 3.

**Figure 3:**
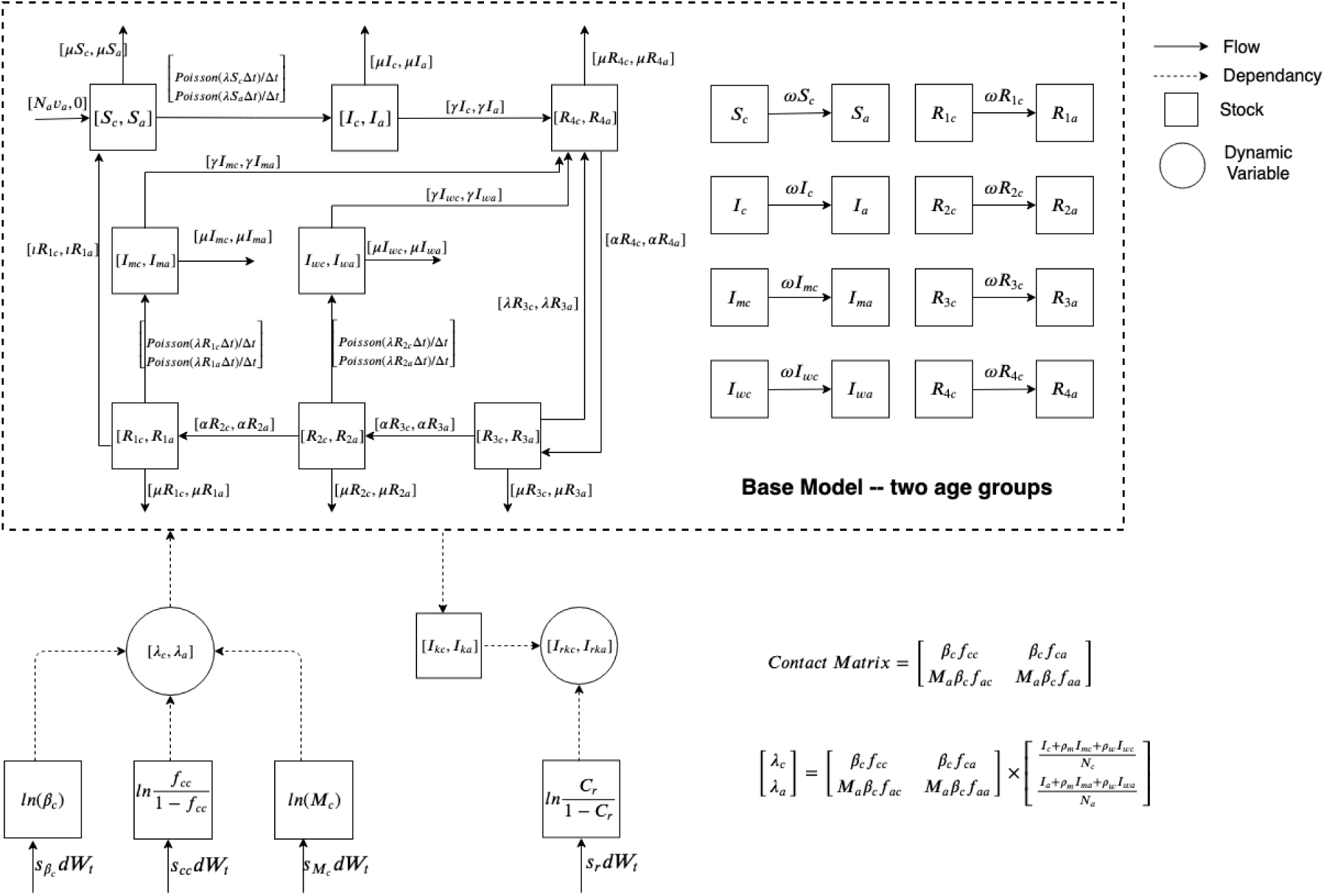
The mathematical structure of the particle filtering age-structured model with two age groups.

As discussed in the aggregate population state space model, we consider occurrence of infections within a given small interval to be characterized by a Poisson process. Then, the flows of new infections incorporated into the model are given by the following equations:

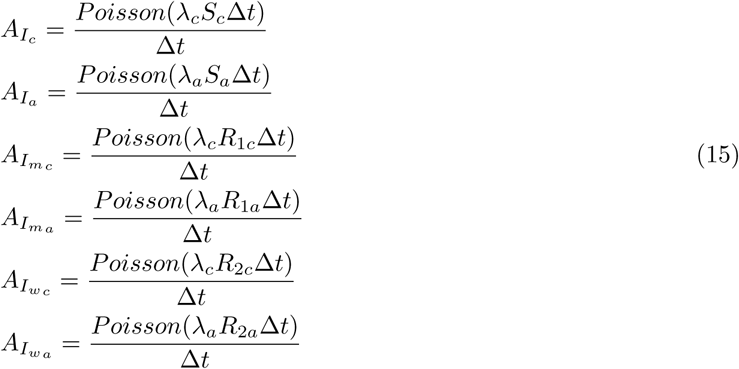

Characterization of the stochastic mixing process between susceptibles and infectives within the stratified model is more involved than the same process in the aggregate population model, due to the need to include both homogeneous mixing within the same age group and heterogeneous mixing amongst different age groups.

In the two-age structured model, we assume that all the differences in transmission from an infected adult vs. an infected child is due to differences in contact rates, and thus that the transmission probability of pertussis (denoted as *p*_*i*_ in the force of infection model of Equation (5) and Equation (6) for age group *i*) are the same between child and adult age groups (i.e., that *p*_*c*_ = *p*_*a*_). Then, according to the general contact matrix model based on mass action introduced previously, we obtain the following equations:

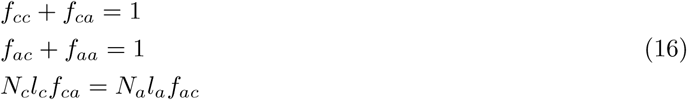

where *l*_*c*_ and *l*_*a*_ are the contact rates of child and adult age groups; *N*_*c*_ and *N*_*a*_ are the total populations of the child and adult age groups; *f*_*ij*_, *i, j* ∈ [*c, a*] indicates the fraction of the contacts of age group *i* occur with age group *j*.

Then, similarly to the aggregate population state space model, we import the parameters – effective contact rates of the child and adult age groups – denoted as *β*_*c*_ and *β*_*a*_, respectively. We know the effective contact rate is the multiplication of the parameter of contact rate and transmission probability. Then, we get *β*_*c*_ = *l*_*c*_*p*_*c*_ and *β*_*a*_ = *l*_*a*_*p*_*a*_. Substituting the equation with *β*_*c*_ and *β*_*a*_ to the Equation (15), we can get [8]:

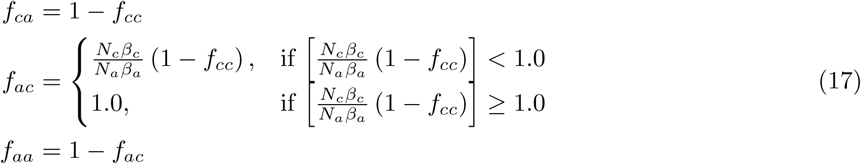

To represent the stochastic characteristics of the mixing process of the two-age group state space model, we allowed three parameters to change with time according to a random walk (with the values of these parameters being estimated as part of model state upon each observation during particle filtering) – the effective contact rate of the child age group *β*_*c*_, the fraction of the contacts of the child age group that occur with the child age group *f*_*cc*_, and the ratio of the adult age group’s effective contact rate (*β*_*a*_) to that of the child age group (*β*_*c*_), denoted as *M*_*a*_. Reflecting the fact that both *β*_*c*_ and *M*_*a*_ vary over the entire range of positive real numbers and *f*_*cc*_ varies in the range of [0, 1], we treat the natural logarithm of each of *β*_*c*_ and *M*_*a*_, as well as the logit of *f*_*cc*_, as undergoing a random walk according to a Wiener Process, and thus undergoing Brownian Motion) [25, 26, 8]. Drawing on notation from the Stratonovich calculus for the random walks involved, we obtain the equations as follows:

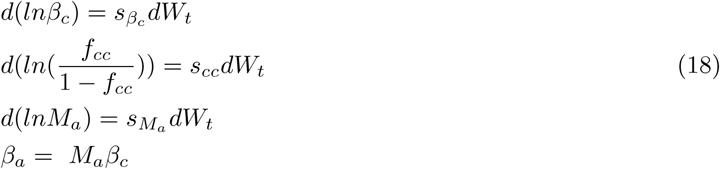

The third stochastic component in the two-age group model relates to calculation of the reported cases of pertussis in the model. As in the aggregate population model, for comparison with reported case counts, we also make use of two convenience states – denoted as *I*_*kc*_ and *I*_*ka*_ – to accumulate pertussis infectious cases from time *k* − 1 to *k* for the child and adult age groups. Moreover, we assume that the pertussis reporting rates of child and adult age groups are the same. Thus, the equation of reporting rate – denoted as *C*_*r*_ – is identical to that in the aggregate model in Equation (12). The mathematical equations characterizing the reporting process are listed as follows:

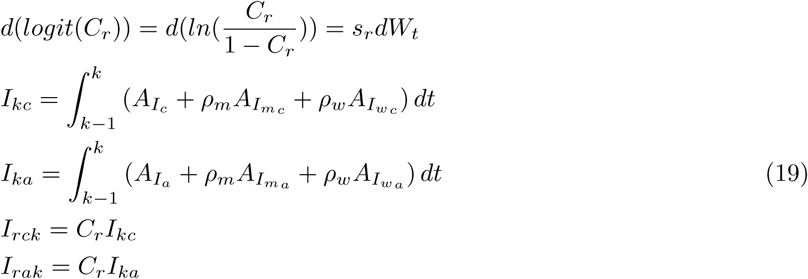

where dynamic variables *I*_*rck*_ and *I*_*rak*_ indicate the reported pertussis cases calculated from the two-age group model.

Finally, the noisy state space model of the two-age group pertussis particle-filtered transmission model is the combination of the base model of the deterministic compartmental epidemiological model in Equations (4) and the adjusted stochastic parts in Equations (15), (18) and (19) (Figure 3). Readers interested in the full mathematical equations of the state space model, values of parameters, and initial states can refer to Appendix D.

##### 32-age group population structure state space models

In this paper, we have explored two pertussis particle filtering models with 32-age group population structure – with the unbalanced contact matrix introduced by [16] (Equation (8)) and re-balanced contact matrix (Equation (9)) – taking the deterministic epidemiological model of Equation (4) with *n* = 32 as the base model. As in the state space models above, we also incorporated three stochastic elements within the 32-age group state space models – the new infectious occurrence process, the contact process between susceptibles and infectives, and the infected case reporting process (Figure 4).

**Figure 4:**
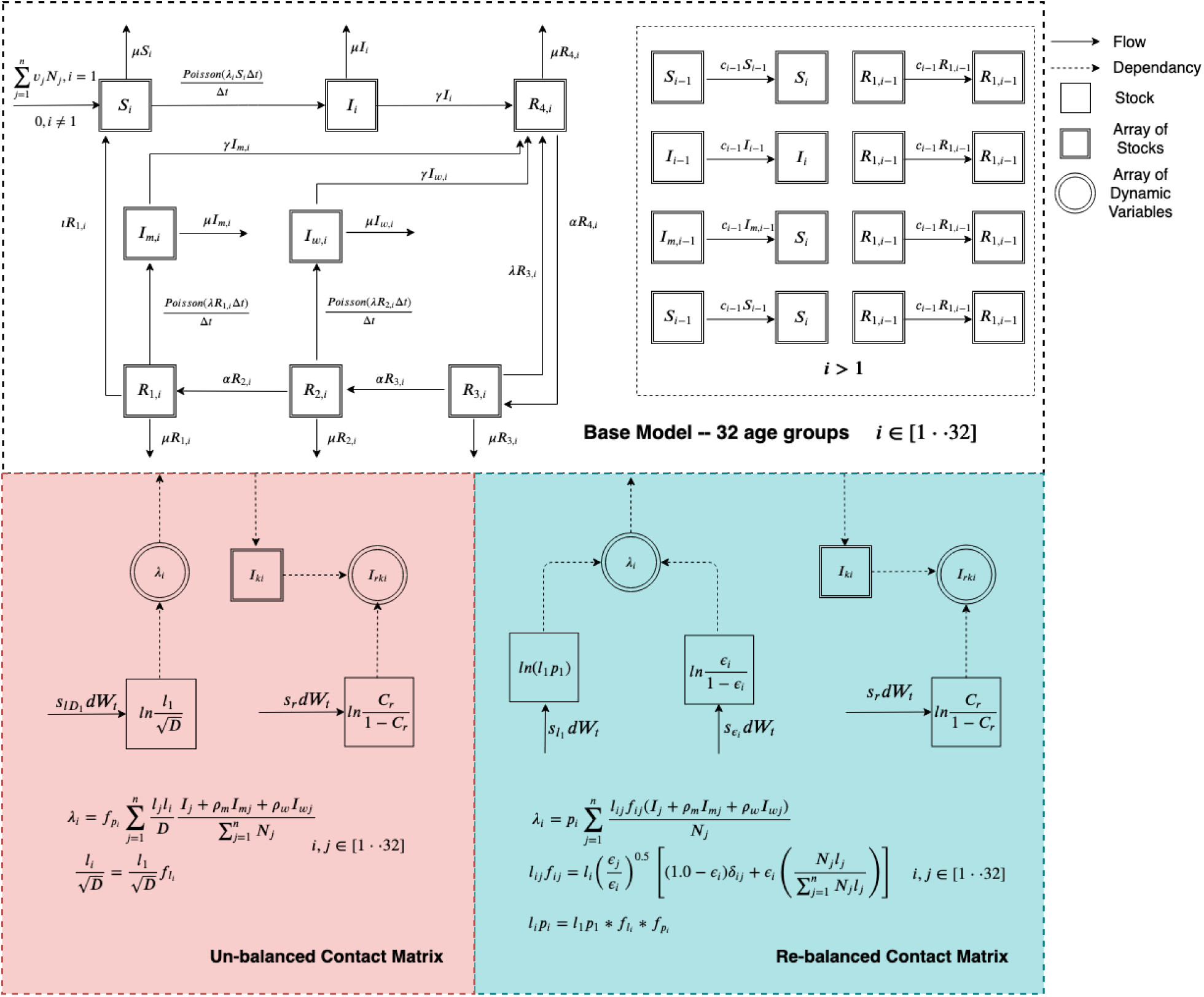
The mathematical structure of the particle filtering age-structured model with 32 age groups.

Similar to the aggregate and two-age group population state space models introduced above, we consider the new infectious individuals occurrence processes follows the Poisson process, and the mathematical equations are listed as follows:

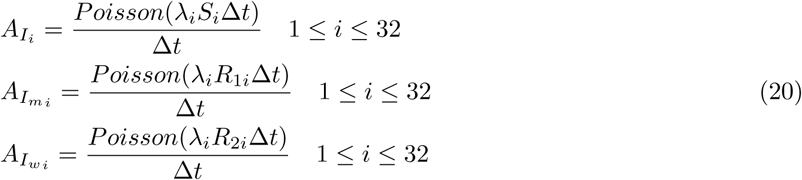

Similarly to those previous models, in the stochastic process of reporting the pertussis cases in the 32-age group state space models, we consider the reporting rate of each age group to be the same, denoted as *C*_*r*_, the logit of *C*_*r*_ undergoing Brownian Motion. The resulting mathematical equations related to the reporting process are listed as follows:

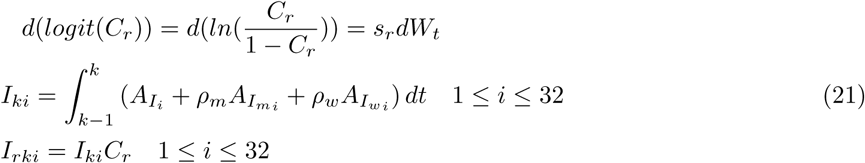

where *I*_*ki*_ represent the new states incorporated into the state space model to capture the accumulative number of pertussis cases from time *k* − 1 to time *k* for age group *i*, and dynamic variable *I*_*rki*_ is the estimated occurrence of reported cases calculated by the state space model for age group *i*.

Then, in characterizing the mixing process between susceptibles and infectives, we separately implement the 32-age group state space model with the un-balanced contact matrix introduced in [16] and re-balanced contact matrix.

In the un-balanced contact matrix method introduced in [16], we consider the parameter of 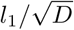 to evolve stochastically in the state space model (i.e., the natural logarithm of 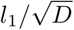 undergoes Brownian Motion). Then, a vector represents how the contact rate of each successive age group compares with that of the first age group; specifically, 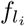 represents the ratio between the contact rate for the first age group and the contact rate of age group *i*. This vector is then used to calculate the parameter of 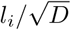 for each age group *i*. 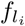 is calculated from the value assumed for contact rate of all age groups, which are taken from [16]. The value of 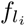 is (1, 6.03, 8.03, 10.03, 12.04, 15.06, 20.08, 28.10, 47.18, 47.18, 47.18, 47.18, 47.18, 25.09, 25.09, 25.09, 25.09, 25.09, 15.06, 15.06, 15.06, 15.06, 15.06, 15.06, 15.06, 15.06, 10.03, 10.03, 5.02, 5.02, 5.02, 5.02). Moreover, another vector 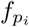 is incorporated to represent the ratio of the transmission probability of pertussis of each age group compared to the first age group in the state space model. The original mathematical model of [16] lacks a dedicated transmission probability parameter. However, one would expect transmission probabilities to different among different age groups. For example, transmission probability from a young child is usually higher than that of the adults due to hygienic disparities. The mathematical equations of the stochastic mixing process are listed as follows:

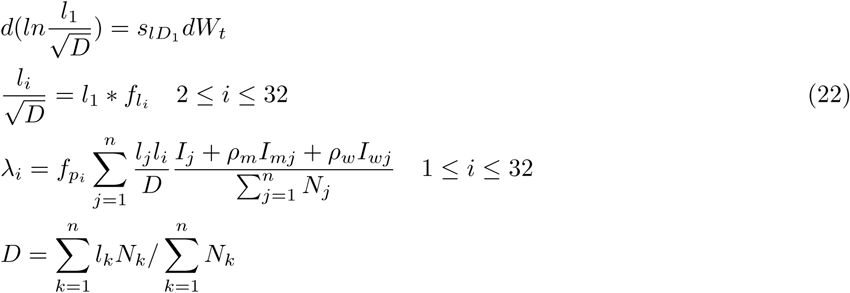

The value of 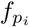 in the 32-age-group models of pertussis in this paper is (1, 1, 1, 1, 1, 1, 1, 1, 1, 1, 1, 1, 1, 1, 1, 1, 1, 1, 0.5, 0.5, 0.5, 0.5, 0.5, 0.05, 0.05, 0.05, 0.05, 0.05, 0.05, 0.05, 0.05, 0.05). We assume that the transmission probability of individuals under 15 years old are the same and is the highest, while the transmission probabilities of individuals in the age groups from 15 to 19 years and over 20 years are half and 1*/*20 compared to the individuals under 15 years, respectively. The population of each age group – which is collected from the age pyramid of Saskatchewan [19] and is assumed to be invariant – is (3349, 3330, 3320, 9950, 19843, 19733, 19647, 19571, 19486, 19394, 19289, 19161, 19002, 18809, 18577, 18318, 18033, 17724, 17386, 17021, 16629, 16218, 15802, 73256, 65935, 117771, 97621, 70964, 44313, 19332, 4377, 387). To let the arrival rate of newborns in each pertussis particle filtering model per unit time (here, month) be the same across all models, the yearly birth rate of the 32-age-group models are assumed as (0, 0, 0, 0, 0, 0, 0, 0, 0, 0, 0, 0, 0, 0, 0, 0, 0, 0, 0.03, 0.03, 0.03, 0.03, 0.03, 0.1, 0.1, 0.1, 0, 0, 0, 0, 0, 0). This is done to ensure the new born population each year are the same in all the models. The values of birth rates are informed from the [20]. Readers interested in a complete characterization of the mathematical equations of 32-age group state space model with an un-balanced contact matrix [16] and the initial values of all states can refer to Appendix D.

In the re-balanced contact matrix method, to represent the stochastic mixing process, we assume that the changes of the logarithm of *l*_1_*p*_1_ (the effective contact rate of the first age group) undergoes a random walk according to a Wiener Process (Brownian Motion) [25, 26, 8]. The logit of the six mixing parameters (*ϵ*_*i*_, 1 ≤ *i* ≤ 6) are similarly treated as evolving according to a Wiener Process. The reason that the total number of mixing parameters is 6, instead of 32 – as might be expected if there are a mixing parameter related to each age group each – lies in the fact that the yearly empirical datasets could only be split into 6 age groups – less than 1 year, 1 to 4 years, 5 to 9 years, 10 to 14 years, 15 to 19 years and 20 years and older, as is characterized in detail below. Finally, the force of infection for the the 32-age group structured pertussis particle filtering model with a re-balanced contact matrix is given as follows:

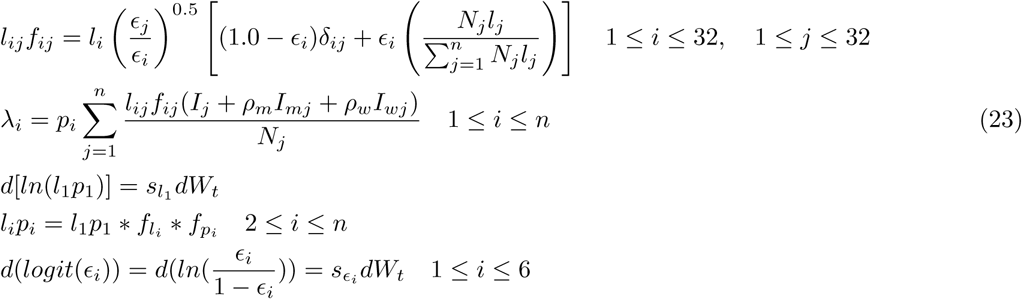

where 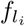 and 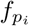 are the ratios of the contact rate and transmission probability between age group *i* and the first age group, respectively. Both the values of 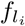 and 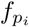 are the same as in the un-blanced contact matrix model. It is notable that we treat the effective contact rate (*l*_*i*_*p*_*i*_) – the multiplication of the contact rate and the transmission probability of age group *i* as a single parameter in this re-balanced model, to be simplify and consistent with the previous models. Thus, we use 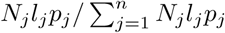 to approximately represent the value of 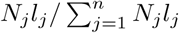 during implementing the model. Readers interested the complete mathematical equations of the 32-age group state space model with a re-balanced contact matrix [16] and the initial values of all states can refer to Appendix D.

#### 2.2.2. Likelihood function

In the condensation method version [23] of the particle filtering method [22], the weight update rule for a particle given a new observation *y*_*k*_ involves multiplying the previous weight by the value of the likelihood function *p*(*y*_*k*_|*x*_*k*_), where the latter represents the probability of observing the empirical data (denoted as *y*_*k*_) given the particle state *x*_*k*_ at time *k*. In this paper, following several past contributions [10, 14, 8, 27], we select the negative binomial distribution as the basis for the likelihood function. We treat the likelihood of observing *y*_*k*_ individuals at time *k* given an estimated count of incident individuals from the model *i*_*k*_ as follows:

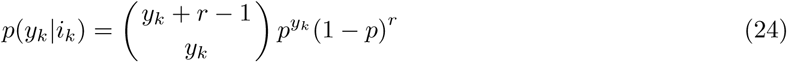

where *y*_*k*_ is the empirical data (reported pertussis cases) at time *k*; *p* = *i*_*k*_*/*(*i*_*k*_ + *r*) represents the probability that a given reported case is in fact a true incident case, and *r* is a dispersion parameter. In all scenarios reported in this paper, the value of *r* is chosen to be 10.

##### Aggregate model

Because the aggregate particle filtering model lacks the capacity to distinguish between individuals with different age groups as necessary to compare to the yearly age-stratified reported values, the measured data for that model consists of a one-dimensional vector giving the reported cases for successive months. The likelihood function in the aggregate model can then correspondingly be calculated by the value of *p*(*y*_*mk*_|*I*_*rk*_), where *y*_*mk*_ is the empirical data as given by the monthly reported measles cases at time *k*, and *I*_*rk*_ is the expected reported cases as calculated by the particle filtering model for each particle.

##### Age structured model

The weight update rule in the age structured model is similar to that in the aggregate model, with the exception of the updates associated with the close of each year. Specifically, we take the likelihood function at the close of the last month (December) of each year as the product of the likelihood functions as formulated for each empirical dataset – including both monthly pertussis reported cases across the whole population and the yearly reported cases related to each of the reported age groups considered. The likelihood formulation of age structured models is as follows:

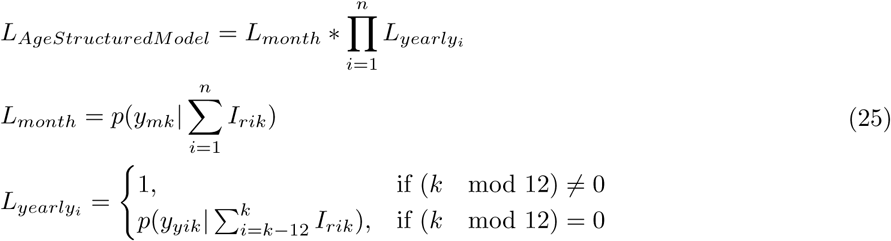

where *L*_*month*_ is the likelihood function based on the monthly empirical data across the total population, 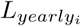 is the likelihood function based on the yearly empirical data for group *i, y*_*yik*_ is the yearly empirical data for age group *i*, and *I*_*rik*_ is the reported pertussis cases of age group *i* at time *k*.

In the two-age group particle filtering model, we have three empirical datasets – the monthly reported pertussis cases across the whole population and yearly reported cases for each of the two age groups (*n* = 2 in Equations (25)). In the 32-age group particle filtering models, we have employed seven empirical datasets – the monthly reported pertussis cases across the whole population and six datasets of yearly reported cases (*n* = 6 in Equations (25)). As noted previously, the yearly empirical datasets could only be split into 6 age groups.

#### 2.2.3. Proposal distribution

The Condensation Algorithm [28, 23] is applied in this project to implement the particle filter model. It is the simplest and most widely used proposal algorithm, making use of the prior as the proposal distribution [23, 22].

### 2.3. Empirical data resources

#### 2.3.1. The surveillance data

This paper benefits from the fact that pertussis is formally classified as a notifiable illness for the mid-western Canadian province of Saskatchewan. Pertussis reporting data for Saskatchewan are used as empirical data for the particle filtering models. These data are public aggregate data obtained from the Government of Saskatchewan’s “Annual Report of Department of Public Health in the Province of Saskatchewan” [20]. This paper employs two categories of datasets drawn from that report – monthly reported cases aggregated across the entire population, and yearly reported cases in each age group. The latter reflects the fact that in the yearly empirical datasets, the annual reported cases are split into different age groups. Within this dataset, age stratification is inconsistent; as a result, the splitting in some years fails to precisely match stratification of the age groups in the models. For these cases, we proportionally split the yearly empirical reported cases into overlapping age groups within the model. Readers interested the detailed introduction of age deviation of the empirical data can refer to Appendix E.

This study employs pertussis reported cases in Saskatchewan specifically during the pre-vaccination era. The monthly empirical data extends from Jan. 1921 to Dec. 1956, with the dataset offering a total of 432 records. Reporting of age-specific data initiated in 1925, and continued through 1956. Every record contains three features – date, reported cases and population size [19]. To make them consistent with the population size of the dynamic model – the average population from 1921 to 1956 (863,545) – the reported cases are normalized to the same population size as the model, as shown in Figure 5, yielding estimated incidence rates rather than incident case counts. It can be readily appreciated that the time series demonstrate the classic patterns of waxing and waning incorporating both stochastic and regular features characteristic of many childhood infectious diseases in the pre-vaccination era.

**Figure 5:**
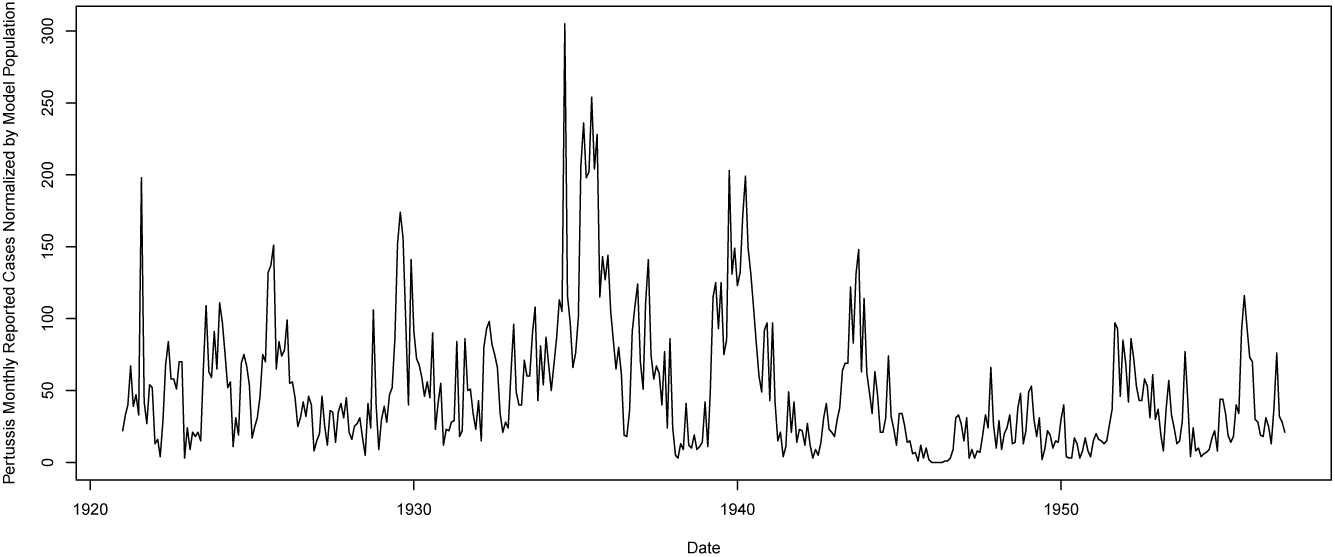
The monthly reported pertussis cases in Saskatchewan from 1921 to 1956 normalized by the population employed in the model (863,545)

#### 2.3.2. The demographic data

The demographic parameters play a significant role in the models, particularly the age structured variants. The parameters related to the population are abstracted from the empirical population of Saskatchewan from 1921 to 1956 [19]. The empirical demographic data indicate that the total population of Saskatchewan does not show drastic fluctuation [19] over the year range from 1921 to 1956. During these years, the empirical population lie in the interval from 757,000 to 932,000. The population of Saskatchewan from 1921 to 1956 of each age is depicted in Figure 6. Thus, we let the model population constantly stay in 863,545, which is the average reported population over the years 1921 to 1956 within the Saskatchewan age pyramid [19]. It bears emphasis that for simplicity, we assumed an equilibrium in the population structure – the total population and population among each age group (in the age-structured models) – remain invariant. Similarly, the model assumes fixed values of the population in each age group, according to the previously noted average population.

**Figure 6:**
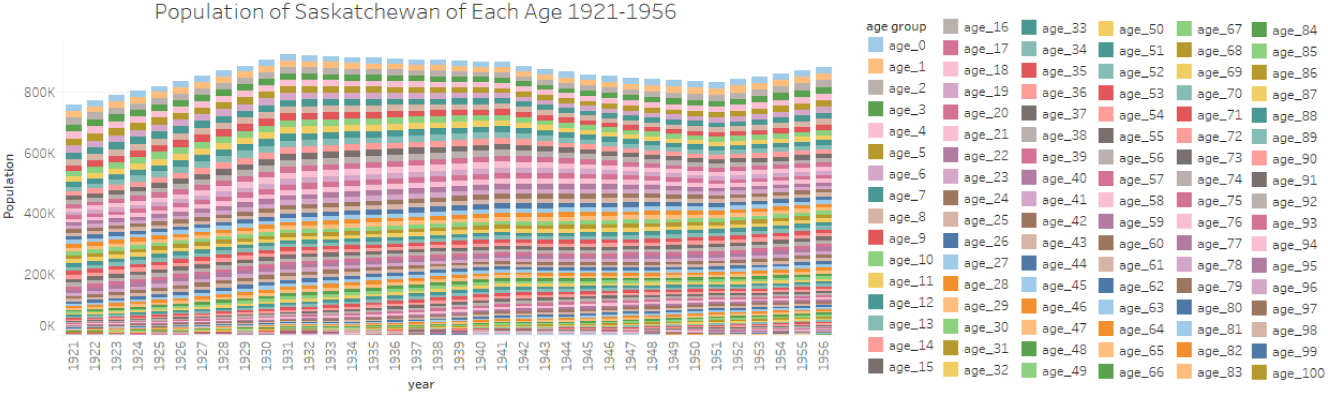
The age-specific and overall population of Saskatchewan from 1921 to 1956.

### 2.4. Introduction of the aggregate population model with calibrated parameters

To evaluate the performance of the particle filtering model when compared to the traditional calibration method, combining with the empirical data, we further constructed a calibration model with the aggregate population using the deterministic epidemiological compartmental model of Equation (1). To be consistent with the particle filtering aggregate model, the parameters and initial values sampled in the particle filtering model are estimated in the calibration model, which are the effective contact rate *β*, reporting rate *C*_*r*_ and the initial value of the stocks of *S, I* and *R*_1_. In this calibrated model, the values of the parameters obtained from calibration against the empirical dataset are listed below. The initial value estimated from the calibration process in class *S, I* and *R*_1_ are 19420, 500, 9960. The value of the effective contact rate (*β*) is 56.692; it bears emphasis that this value incorporates both a rate of contact and the probability of transmission. The calibrated value of the reporting rate of pertussis is 0.01. The other parameters are the same as the particle filtering models.

### 2.5. Classifying outbreak occurrence

The pertussis particle filtering models – combining the particle filtering algorithm and the compartmental models with empirical data – are capable to estimate and predicting the full (continuous) model state over time. Moreover, in this paper, on the basis of having particle filtered up to a certain month, we further perform classification outbreak (outbreak vs. non-outbreak) analysis based on the predicted results (in the next time unit – month) of the particle filtered models. Referred from our previous contribution [8], the function mapping from the continuous predicted results of particle filtering models – predicted reported pertussis cases in the next month – to dichotomous categories of outbreak and non-outbreak can be represented as follows [23, 8]:

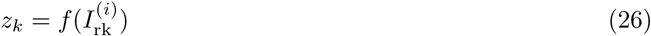

where 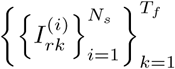 indicates the matrix of reported cases of pertussis predicted by the particle filtering model of particle *i* (1 ≤ *i* ≤ *N*_*s*_) at time *k* (1 ≤ *k* ≤ *T*_*f*_). *T*_*f*_ is the total running time of the model.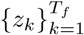 is the vector of dichotomous predicted classes – *z*_*k*_ *∈* {0, 1}, where 0 indicates non-outbreak, and 1 indicates outbreak. The value *I*_rk_ is generated by the particle filtering models. Specifically, *I*_*rk*_ equals 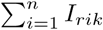 (where *I*_*rik*_ is the reported pertussis cases of age group *i* at the time *k*) in the particle filtering models introduced above.

Two processes are then used to perform classification analysis of the results from the particle filtering models [8]. In the first process, we define a threshold (*θ*) – mean plus 1.5 times the standard deviation of the empirical monthly reported cases, above which that particle is considered as positing an outbreak. In the second process, we define a threshold of the fraction (*θ*_*k*_) of particles required to posit an outbreak at time *k* for us to consider there as being an outbreak. Then, the vector determining whether there is an outbreak of measles in each month – *z*_*k*_ – is calculated. We further denote 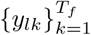 as the binary empirical vector of whether a pertussis outbreak indeed obtained at time *k, y*_*lk*_ ∈ {0, 1}. The calculation method of *y*_*lk*_ is similar to that of each particle. If the count of measles reported cases is greater or equal to the threshold *θ*, the related element in vector *y*_*lk*_ is labeled to be outbreak (the value is 1). Otherwise, a non-outbreak is assumed (the value is 0).

Finally, to summarize the performance of the classifier, we employ as a metric the area under the Receiver Operating Characteristic (ROC) curve. Readers interested in additional detail are referred to our previous contribution utilizing a comparable methodology for measles [8].

## 3. Results

### 3.1. Results of models incorporating empirical datasets across all timeframe

Recall that to explore the predictive performance of particle filtering in different compartmental pertussis models, four distinct particle filtering models have been built in this research – the aggregate particle filtering pertussis model (denoted as *PF*_*aggregate*_), the age-structured particle filtering model with 2 age groups (denoted as *PF*_*age_*2_), the age-structured particle filtering model with 32 age groups with the original Hethcote contact matrix (denoted as *PF*_*age_*32*_Hethcote*_), and the age-structured particle filtering model with 32 age groups with the re-balanced contact matrix (denoted as *PF*_*age_*32*_rebalanced*_). In each of the four particle filtering models, 3000 particles are used in the particle filtering algorithm; for clarity in exposition, we sampled the same number when generating the plots of the 2D histogram and for calculating the discrepancy. To compare the accuracy of a particle filtered model against that of a traditional model of pertussis calibrated against comparable data, we have further built a calibrated model of the aggregate population, henceforth denoted *Calibrated*.

By comparing the discrepancy – the root mean square error (RMSE) between the model results and the empirical data – associated with each model, we sought to identify the model offering the greatest predictive validity. We then use the most favorable model to perform prediction and intervention analysis.

Each of the five particle filtering models was run 5 times (the random seed generated from the same set). Shown here are the average and 95% confidence intervals (in parentheses) of the mean discrepancy for each model variant.

To assess model results, each of the four particle filtering models was run 5 times with random seeds generated from the same set. We calculated the average and 95% confidence intervals of the mean discrepancy. Table 1 displays the average discrepancies of the four pertussis particle filtering models and the calibrated deterministic pertussis compartmental model, where the discrepancy considers the entire timeframe. These results suggest that particle filtering models significantly improve the predictive accuracy beyond what is achieved via calibration. It is notable that both the calibrated deterministic model and the aggregate particle filtering model only offer monthly average discrepancy, because the yearly observations are stratified by age, but age stratification absent in both such models. Table 1 indicates that the particle filtering models are significantly more accurate than the calibrated model – the average discrepancies of the particle filtering models are significantly lower than those for the calibrated deterministic model. Moreover, although the monthly average discrepancies among the four particle filtering models with different population structure and contact matrix structure are quite close, the particle filtering models *PF*_*age_*2_ and *PF*_*age_*32*_rebalanced*_ exhibit smaller average discrepancies. With respect to the yearly average discrepancies, Table 1 shows that the age-structured model with two age groups offers better predictive performance than the model with 32 age groups; as noted, the aggregate model lacks the age stratification required to calculate yearly discrepancies. It is notable that the total number of the yearly empirical datasets against which the calibration is assessed is different between the age-structured models with 2 age groups (which is compared with 2 yearly empirical datasets) and that with with 32 age groups (which is compared with 6 empirical yearly datasets). The yearly average discrepancies listed in Table 1 are the sum of the average discrepancy across each empirical dataset. Thus, this difference may contribute to the result that the yearly average discrepancies of the model with 32 age groups are greater than the model with 2 age groups; at the same time, this effect will tend to be limited by the fact that both the model and the empirical values will tend to have smaller counts when applied to a greater number of age groups, yielding a smaller per-age-group discrepancy. On balance, we chose to employ the particle filtering model with two age groups as the minimum average discrepancy model to explore the performance of pertussis outbreak prediction.

**Table 1:** Comparison of the average discrepancy (RMSE) for the calibrated model and all four particle filtered pertussis models, considering empirical data across all observation points; parentheses give the 95% confidence intervals.

| Model | Monthly | Yearly in Month | Total |
| --- | --- | --- | --- |
| *Calibrated* | 34.2 | NONE | NONE |
| $PF_{aggregate}$ | 20.9 (20.0, 21.9) | NONE | NONE |
| $PF_{age\_2}$ | 19.9 (18.8, 21.0) | 21.0 (19.2, 22.7) | 40.9 (38.1, 43.6) |
| $PF_{age\_32\_Hethcote}$ | 20.6 (20.1, 21.2) | 25.8 (23.0, 28.6) | 46.4 (43.1, 49.7) |
| $PF_{age\_32\_rebalanced}$ | 19.8 (19.5, 20.1) | 28.1 (24.2, 31.9) | 47.9 (43.9, 51.9) |

Figure 7 shows a boxplot of the distribution of discrepancies among the calibrated model and the four particle filtering models, where a given box in the boxplot summarizes monthly discrepancy estimates for a given model, where those discrepancies are considered over different points in time. Each of the particle filtering models was run 5 times (with the random seed being generated from same set). Then the average monthly and yearly discrepancy among these five runs at each observation time between the particle filtering models and the empirical data are recorded for the boxplot. Both the monthly and yearly (adjusted to units of one Month by dividing by 12) distributions of the discrepancies of each of the age structured models are plotted in Figure 7. This boxplot also indicates that when considered over time, the the discrepancies of all the particle faltering models tend to be smaller than for the calibrated model, although there are similar median discrepancy values. More notable yet is the fact that the discrepancies associated with the calibrated model are significantly more variable than those for the particle filtered models. This suggests that particle filtering improves the consistency of the model’s match against empirical data, when compared to a traditional deterministic model with calibrated parameters. Finally, it bears note that the datasets of the discrepancy of the model *PF*_*age_*2_ have a particularly narrow distribution, especially when judged in terms of yearly discrepancy.

**Figure 7:**
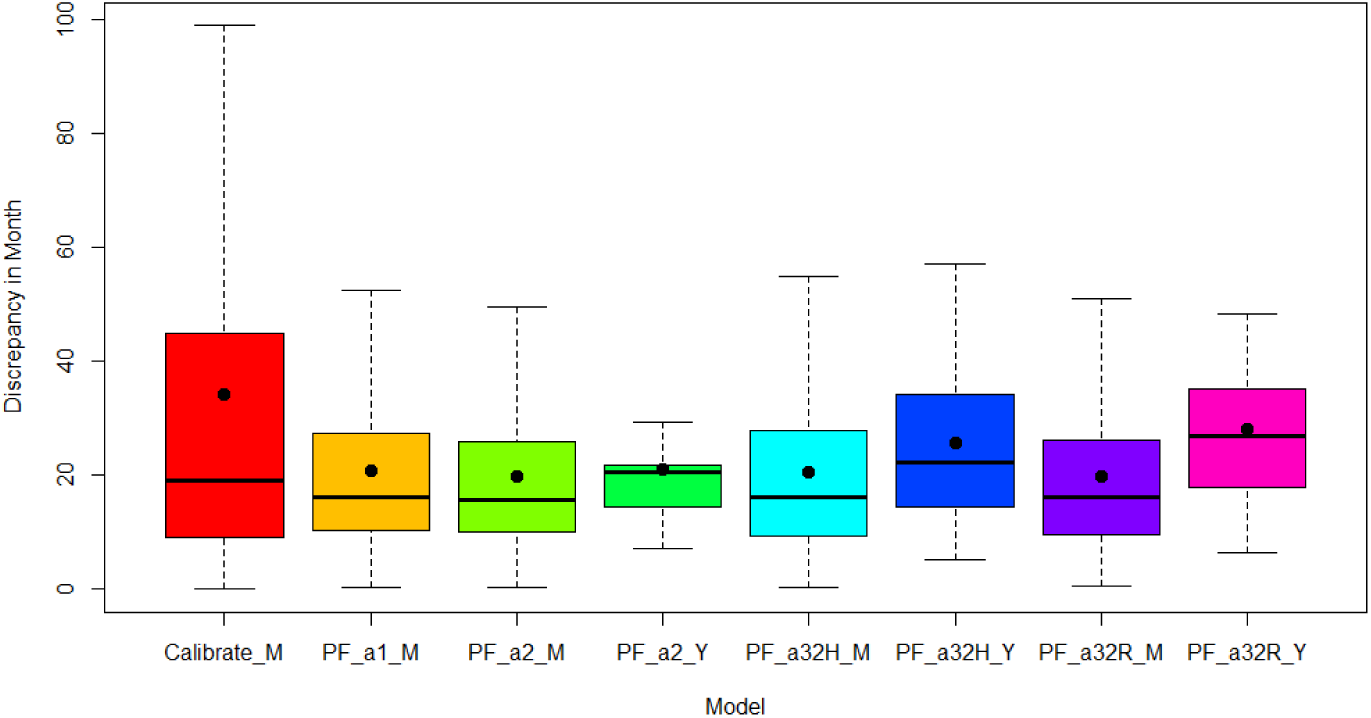
Boxplot of monthly and yearly discrepancy of all models at monthly observation points, considering empirical data across all observation points. “Calibrate” indicates the calibration model with aggregate population structure; “PF a1” indicates the particle filtering model with aggregate population structure; “PF a2” indicates the particle filtering model with 2 age groups; “PF a32H” indicates the particle filtering model with 32 age groups and the contact matrix introduced in [16]; “PF a32R” indicates the particle filtering model with 32 age groups and the re-balanced contact matrix. “M” indicates the discrepancy of the model comparing model-based monthly results with the monthly empirical data – the pertussis reported cases among all population; “Y” indicates the sum of discrepancy (of each age group) of the models comparing model-based yearly results with the yearly empirical data – the pertussis reported cases classified into age groups and having adjusted the unit to Month by dividing by 12. It is also notable that the dot in the boxplot indicates the mean value, while the horizontal line indicates the median value.

Figure 8 compares the output of the calibration model and the empirical data. It indicates that the deterministic model even with parameters calibrated against the entire scope of data encounters difficulties in tracking oscillations associated with waning and waxing of pertussis almost across the entire model time horizon, reflecting the approach of the deterministic model towards a stable equilibrium. These results indicate that the particle filtering models considered here can not only decrease the discrepancy between model results and the empirical data, but can further track the oscillation of outbreaks of pertussis.

**Figure 8:**
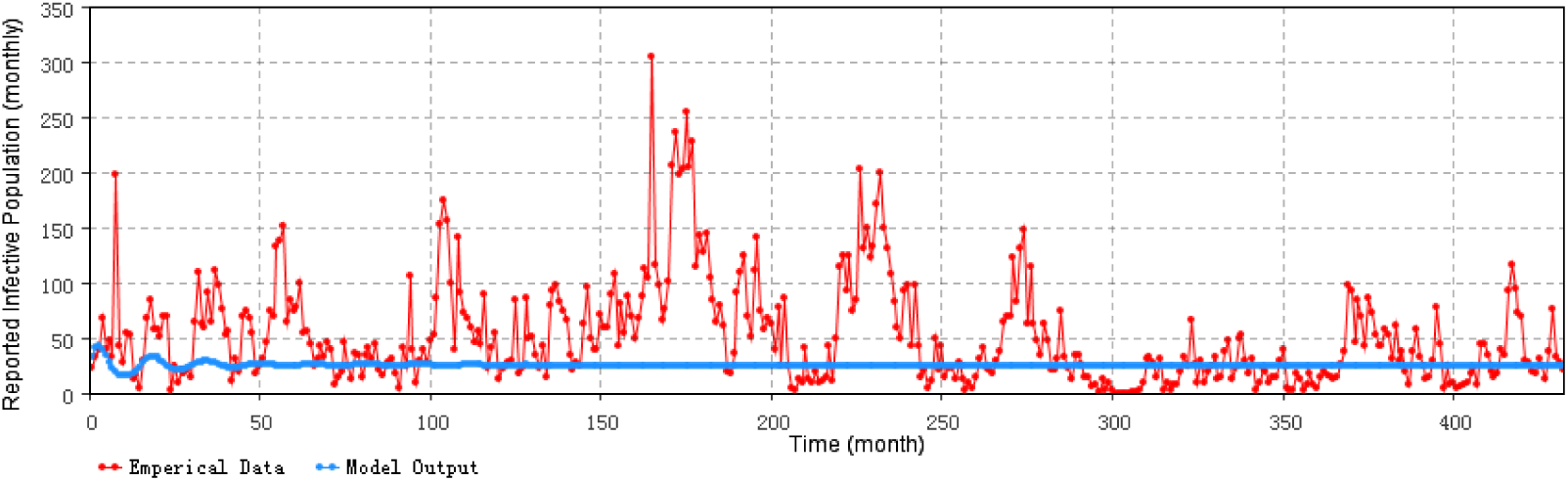
Reported pertussis cases predicted by the calibration model (monthly).

Taken together, the results shown in Figure 7 and Figure 8 suggest that incorporating particle filtering in the compartmental model of pertussis could enhance simulation accuracy and support more accurate outbreak tracking.

Figure 9 presents the posterior results of the pertussis particle filtering model with aggregate population structure over the entire timeframe. For this diagram at time *t*, the results of the particle filtering model at time *t* are sampled according to the weight of all particles following the update to those weights resulting from incorporating the empirical data from time *t*. Those time-specific values are then plotted; the values of empirical data points are shown in red, while the sampled posterior distribution of particle filtering model are shown in blue. The blue color saturation indicates the relative density of sampled points within a given 2D bin. Figure 9 demonstrates that most of the empirical data points are located in or near the high density region of the posterior distribution of the particle filtering model. The results shown in that figure further indicate that the particle filtering model has the capability to track the outbreak of pertussis over time, especially compared with the calibrated model whose results are shown in Figure 8. It bears emphasis that the particle filtered results can follow the patterns of empirical data as they arrive; this capacity to update its estimate of model state – both latent and observed – in with new arriving data is central to the function of particle filtering. By contrast, calibration lacks a means of updating the estimate of the model state over time, and is instead relegated to estimating parameter values, rather than the values of the state at varying points in time.

**Figure 9:**
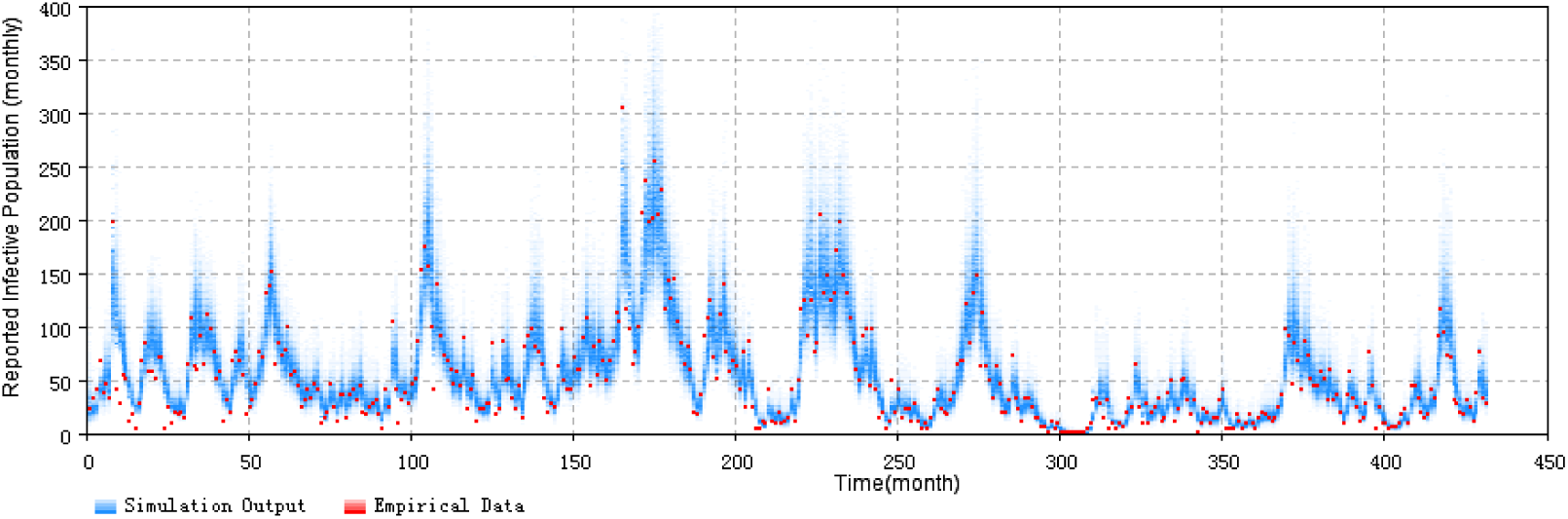
2D histogram posterior result over the total timeframe for the aggregate particle filtering pertussis model. The posterior result is sampled following weight updates in light of observations of empirical data arriving at each unit time.

Figure 10 shows the prior results of the pertussis particle filtering model with aggregate population structure for the entire timeframe. For the prior diagram, the results are sampled before the weight update step triggered by arrival of an empirical data point. Compared with the posterior results shown in Figure 9, the prior values of sampled particles of Figure 10 are distributed over a wider range. This difference in dispersion indicates that the weight update process of particle filtering algorithm in this paper has the capability to use an empirical datum to concentrate the distribution of particles in the state space of the particle filtering model into a tighter range offering greater consistency with the empirical datum.

**Figure 10:**
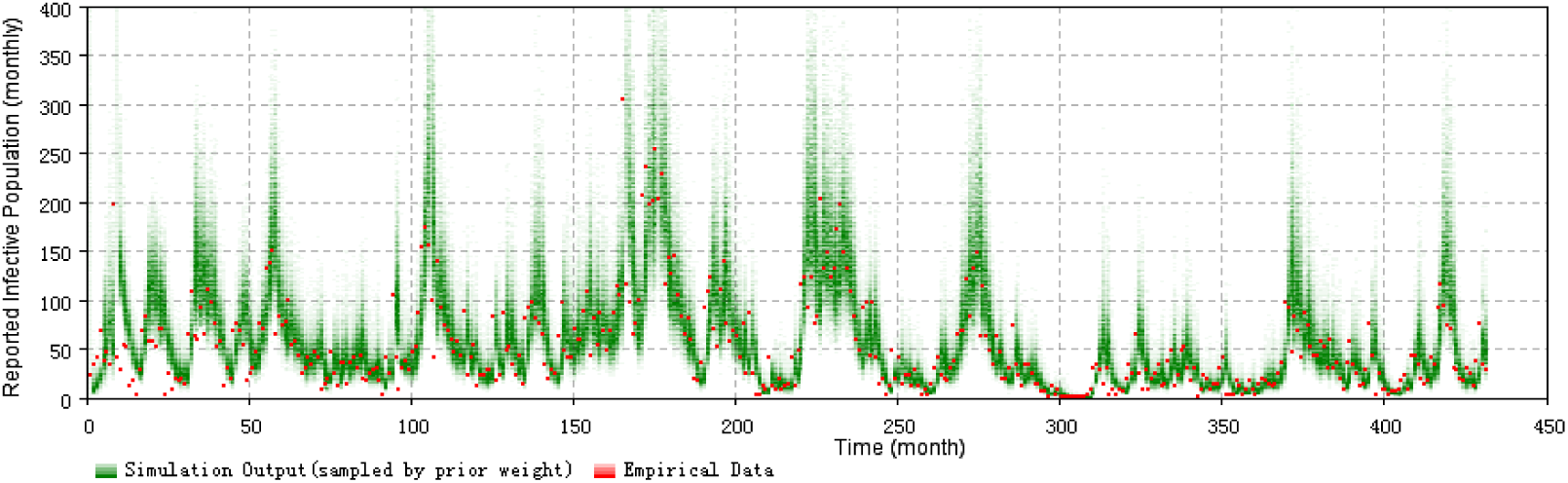
2D histogram prior result over the total timeframe for the aggregate particle filtering pertussis model. The prior result is sampled before the weight updates in light of observations of empirical data arriving at each unit time.

Figure 11 displays the 2D histogram plots comparing both the monthly and yearly empirical datasets (on the one hand) with the distributions of samples from the posterior distribution of incident cases from the age structured pertussis particle filtering model containing 2 age groups (denoted as *PF*_*age_*2_) (on the other). This figure demonstrates that the model *PF*_*age_*2_ is capable of tracking and simulating outbreaks of pertussis, as evidenced by the fact that most of the monthly and yearly empirical data (shown in the red dashes) in each month are located in or near the high density region of the sampled distribution of the particle filtering model (shown in blue in the 2D histogram plots).

**Figure 11:**
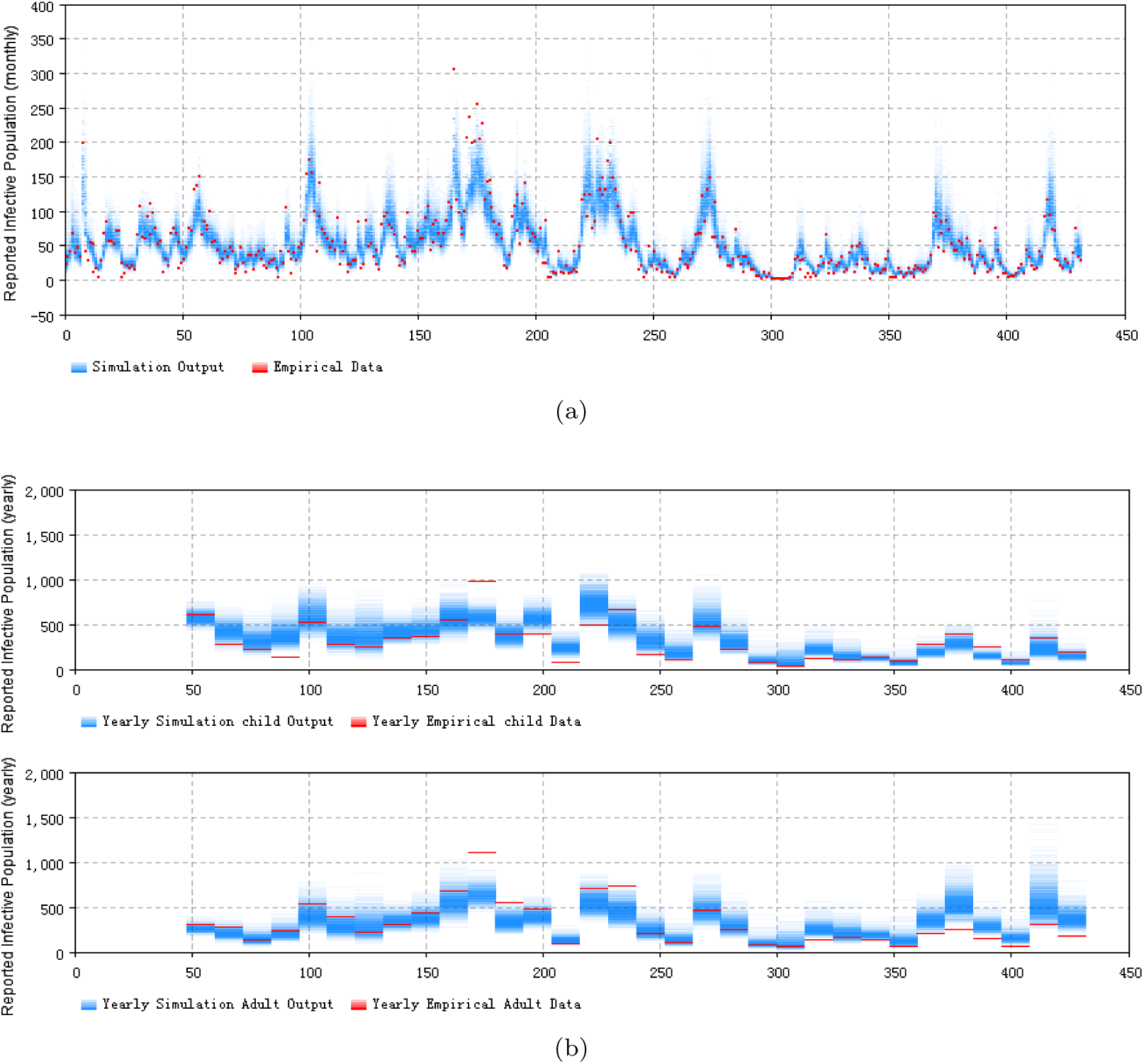
2D histogram posterior result over the total timeframe of the two-age stratified pertussis model. (a) the monthly particle filtering result summed over the entire population. (b) the yearly particle filtering result for the child (top) and adult (bottom) age groups.

Figure 12 displays the 2D histogram plots comparing both the monthly and yearly empirical datasets (on the one hand) with the sampled posterior distribution of incident cases from the age structured pertussis particle filtering model with 32 age groups and the Hethcote contact matrix (denoted as *PF*_*age_*32*_Hethcote*_) (on the other). It is notable that the total number of the yearly empirical datasets employed is 6. This figure also demonstrates that the model *PF*_*age_*32*_Hethcote*_ is capable of tracking and simulating the outbreaks of pertussis, as reflected in the fact that most of the monthly and yearly empirical data for each observation point are located in or near the high density area of the results of the particle filtering model.

**Figure 12:**
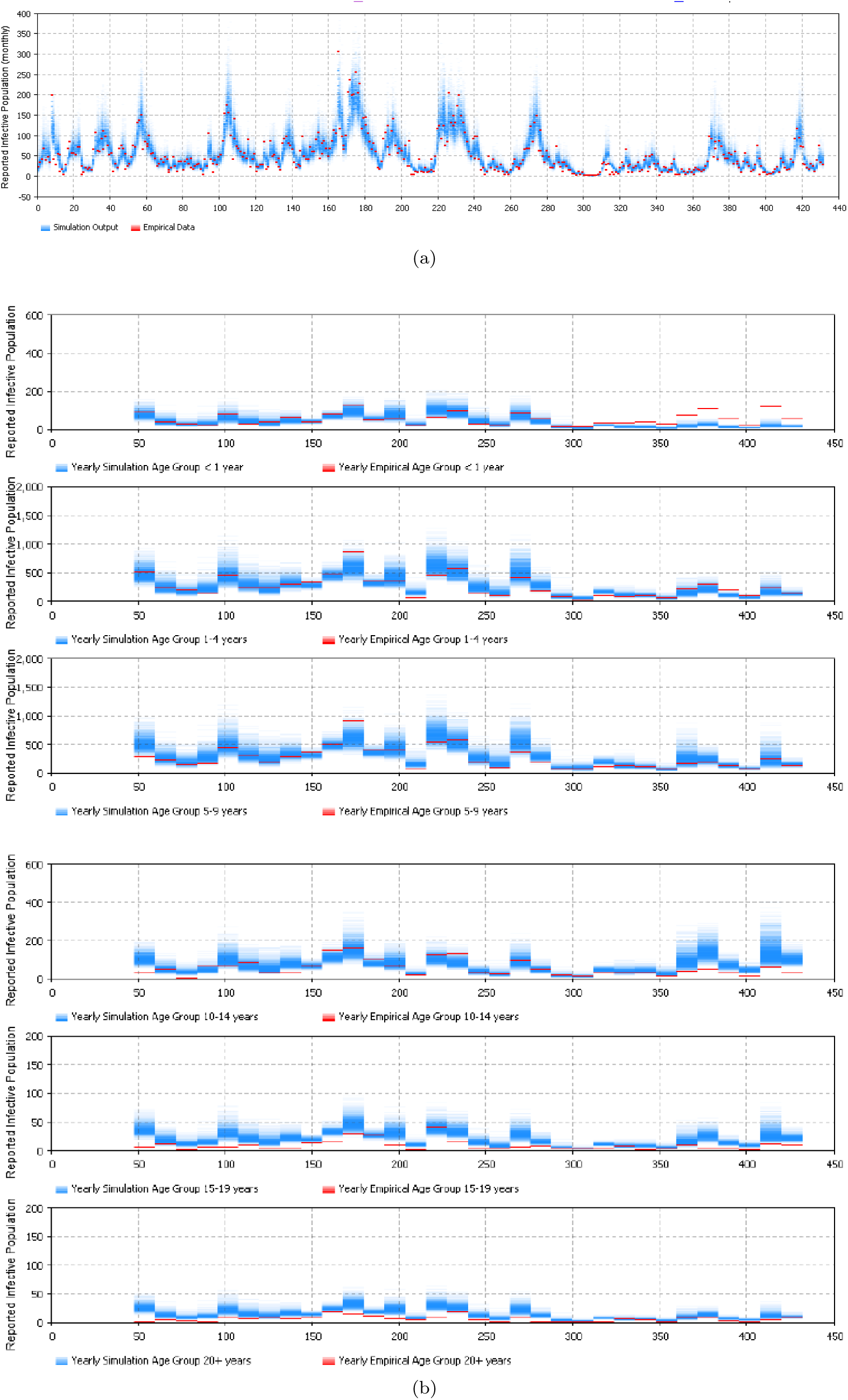
2D histogram posterior result over the total timeframe of the age structured model of 32 age groups with the Hethcote contact matrix. (a) the monthly particle filtering result summed over the entire population. (b) the yearly particle filtering results of each age group of empirical datasets; age groups are successively older from top to bottom.

Figure 9, Figure 11 and Figure 12 represent the 2D histogram posterior result of all the particle filtering models, except for the age-structured model of 32 age groups with a re-balanced contact matrix. Results are omitted for this final model as they are highly similar to those for the 32-age-group model using the Hethcote contact matrix, which is itself shown in Figure 12. The 2D histogram plots shown indicate that both the age-structured particle filtering models and the aggregate population particle filtering model have the capability to closely track the outbreak pattern of pertussis. The results of the models could match the empirical datasets quite well, including both monthly empirical dataset and yearly empirical datasets. In contrast to the calibrated model whose results are shown in Figure 8, the particle filtering models are capable of localizing the model’s prediction of empirical data near the empirical data, as achieved by concentrating the distribution of particles across the underlying state space. Although the results in Table 1, Figure 7 (for discrepancy), Figure 9, Figure 11, and Figure 12 (for posterior distribution) suggest that all four pertussis particle filtering models are capable of tracking and estimating the pertussis outbreaks, in the interest of brevity of exposition, we selected the minimum discrepancy model – the age-structured particle filtering pertussis model with 2 age groups – to perform the prediction and intervention analysis below.

### 3.2. Prediction of outbreaks with the minimum discrepancy model

To assess the predictive capacity of the pertussis particle filtering models in anticipating outbreaks, we performed out-of-sample prediction experiments. Informally, each such experiment examines the capacity of the model to project results into the future, having considered data only to some “current” time. That is, the model is particle filtered so as to incorporate data only to up to – but not including – a “Prediction Start Time” (*T*\*), and then begins projecting (predicting) forward, starting at *T*\*. More specifically, in this process, the weights of particles will cease updating in response to observations at time *T*\*; following that point, all of the particles run without new empirical data being considered. In this paper, all of the prediction experiments are run for 4 years following the “Prediction Start Time” *T* *. To evaluate the predictive capacity of the model, we examined the effects of changing the prediction start time *T*\* so as to pose different archetypal types of prediction challenges. It is notable that the minimum discrepancy model – the age structured model with 2 age groups where the child age group represents children in the first 5 years of their life, and incorporating both the monthly and yearly empirical datasets, as identified in the previous section – is employed to perform all of these experiments.

1. Prediction started from the first or second time points of an outbreak.
2. Prediction started before the next outbreak.
3. Prediction started from the peak of an outbreak.
4. Prediction started from the end of an outbreak.

Figures 13–16 display the prediction results of these situations with respect to the monthly 2D histogram of population-wide reported case counts. In the 2D histogram plots of Figures 13–16, the empirical data having been considered in the particle filtering process (i.e., incorporated in training the models) are shown in red, while the empirical data considered in the particle filtering process (and only displayed to compare with model results) are shown in black. The vertical straight line labels the “Prediction Start Time” (*T*\*) of each experiment.

**Figure 13:**
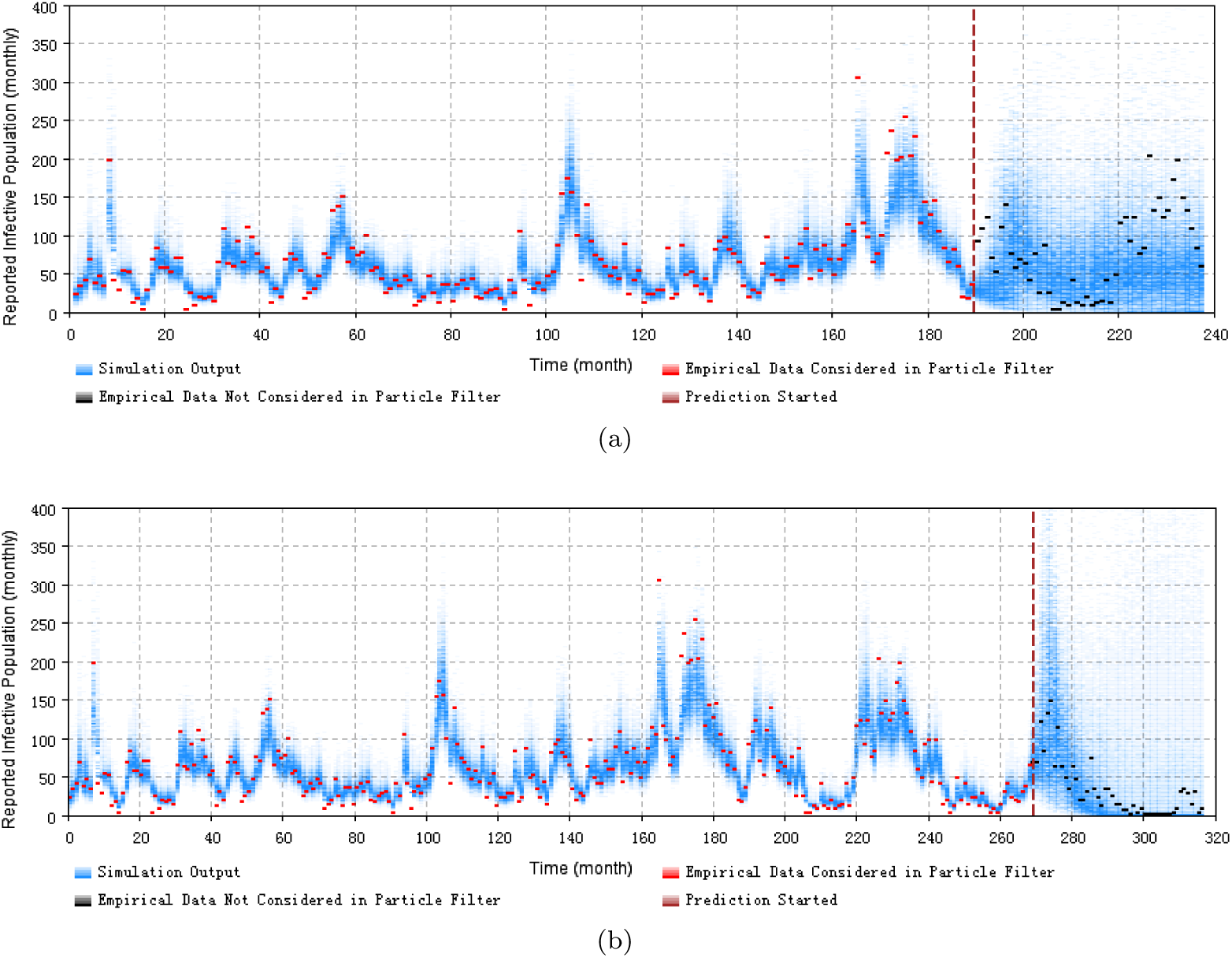
2D histogram depicting prediction using the minimum discrepancy model from the first or second time points of an outbreak. (a) prediction from month 190. (b) prediction from month 269.

**Figure 14:**
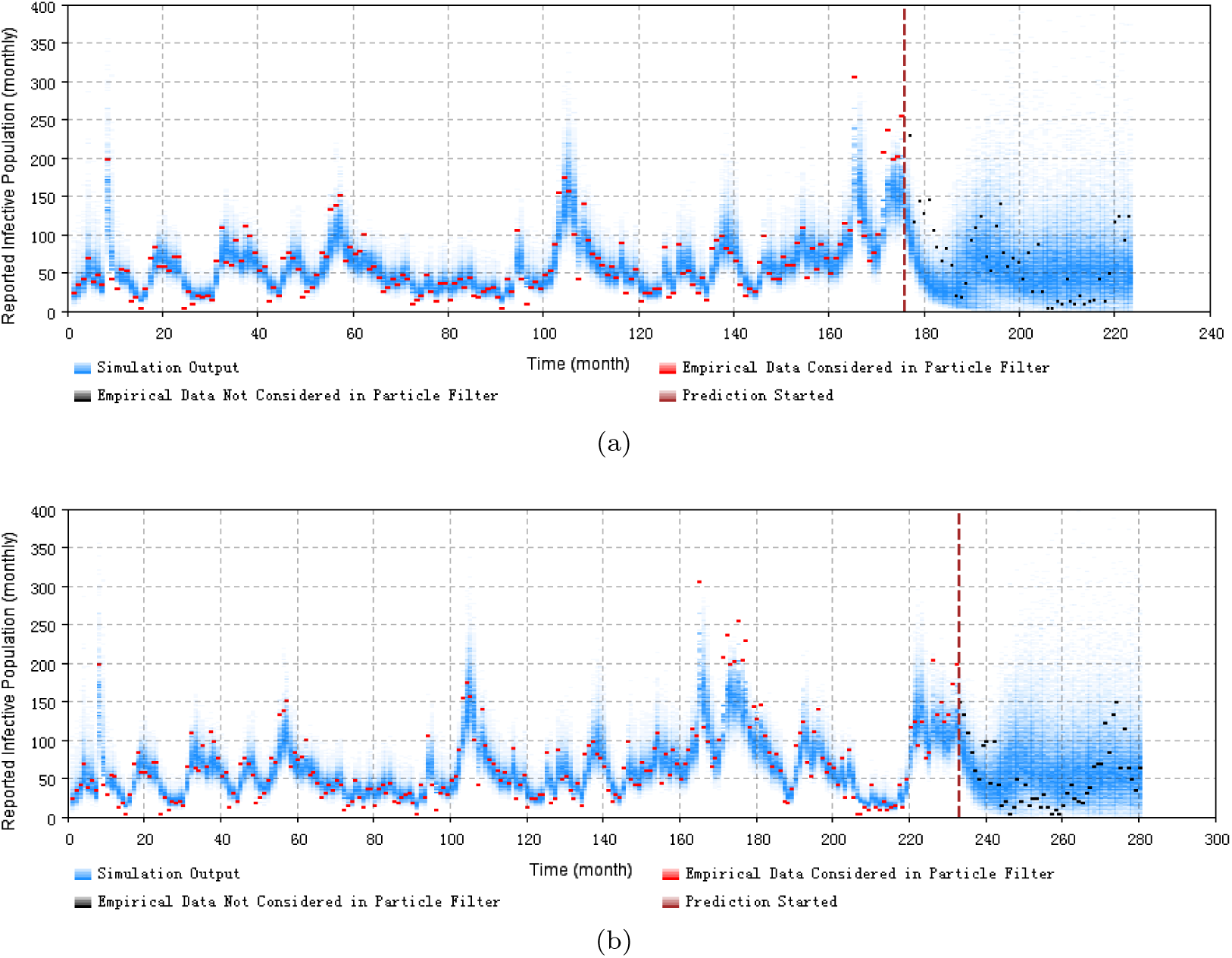
2D histogram depicting prediction using the minimum discrepancy model from the peak of an outbreak. (a) prediction from month 176. (b) prediction from month 233.

**Figure 15:**
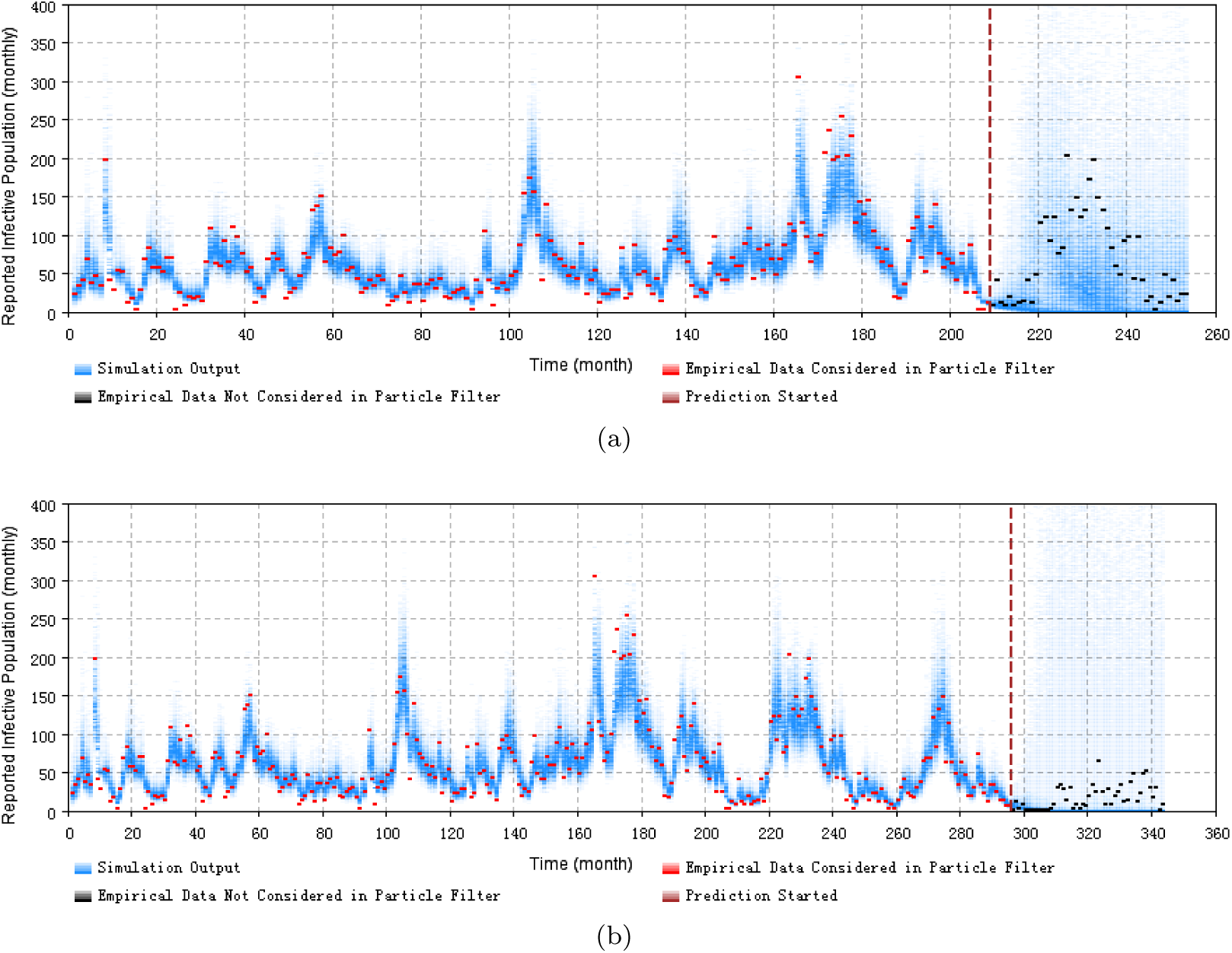
2D histogram depicting prediction using the minimum discrepancy model from the end of an outbreak. (a) prediction from month 209. (b) prediction from month 296.

**Figure 16:**
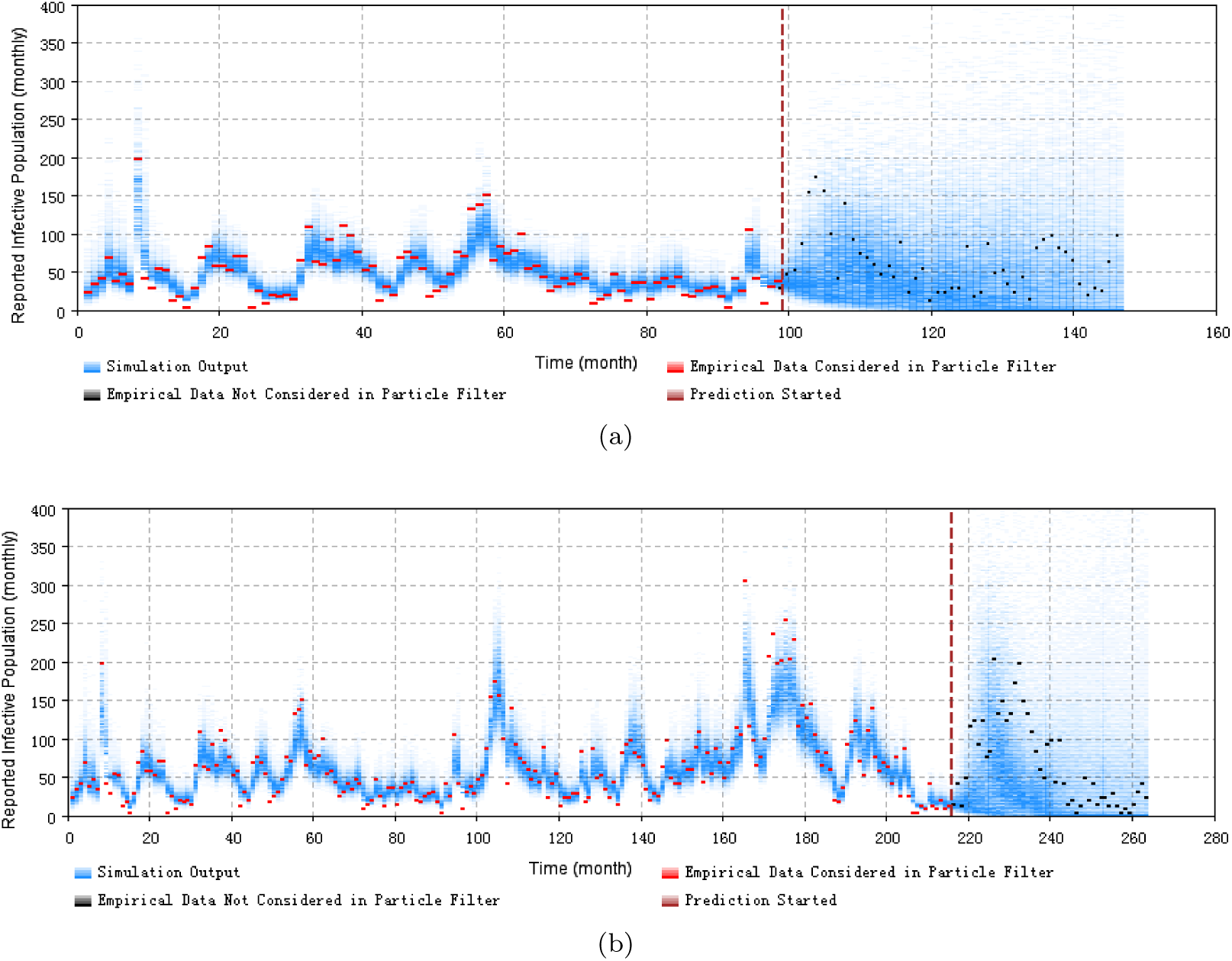
2D histogram depicting prediction using the minimum discrepancy model prior to the next outbreak. (a) prediction from month 99. (b) prediction from month 216.

These prediction results suggest that the pertussis particle filter model offers the capacity to probabilistically anticipate pertussis dynamics with a fair degree of accuracy over a year or so. From the 2D histogram plots, empirical data lying in the projection interval after the prediction start time – and thus not considered by the particle filtering machinery – mostly lie within the high-density range of the particles. Reflecting the fact an ability to accurately anticipate a high likelihood of a coming outbreak could offer substantial value for informing public health agencies with accurate predictions of the anticipated evolution of pertussis over coming months, the next section formally evaluates the performance a simple classifier as to whether the next month will be subject to an outbreak or not, where that classifier uses a very simple prediction scheme constructed atop the particle filter model.

### 3.3. Prediction of classifying outbreak occurrence of the minimal discrepancy model

Beyond assessing the use of particle filtering models for predicting forward pertussis transmission more generally, we also used the lowest discrepancy particle filter pertussis model (*PF*_*age_*2_) to dichotomously predict occurrence of a pertussis outbreak within the next month.

Figure 17 displays an evaluation of the predictive performance in the form of an ROC curve. The Area Under the Curve (AUC) of the ROC curve is 0.913, suggesting that it is possible to achieve both high specificity and high sensitivity. Figure 18 shows the boxplot of residuals (difference between predicted model result and empirical data) of sampled particles (by weight) at each time point where empirical data comes in (each month). Two points bear emphasis. Firstly, these results depict prior model predictions – that is, those predicted by the model before the new data is observed. Secondly, Figure 18 excludes the first 10 months (empirical data points) of the time horizon, during which the particle filtering model is not stable enough due to insufficient incorporation of empirical data. Figure 18 indicates that for results of the next time point (month in this paper), the prior prediction of the particle filtering model are quite close to those of the empirical data – although the empirical data at each predicted time point are not yet incorporated to ground the model.

**Figure 17:**
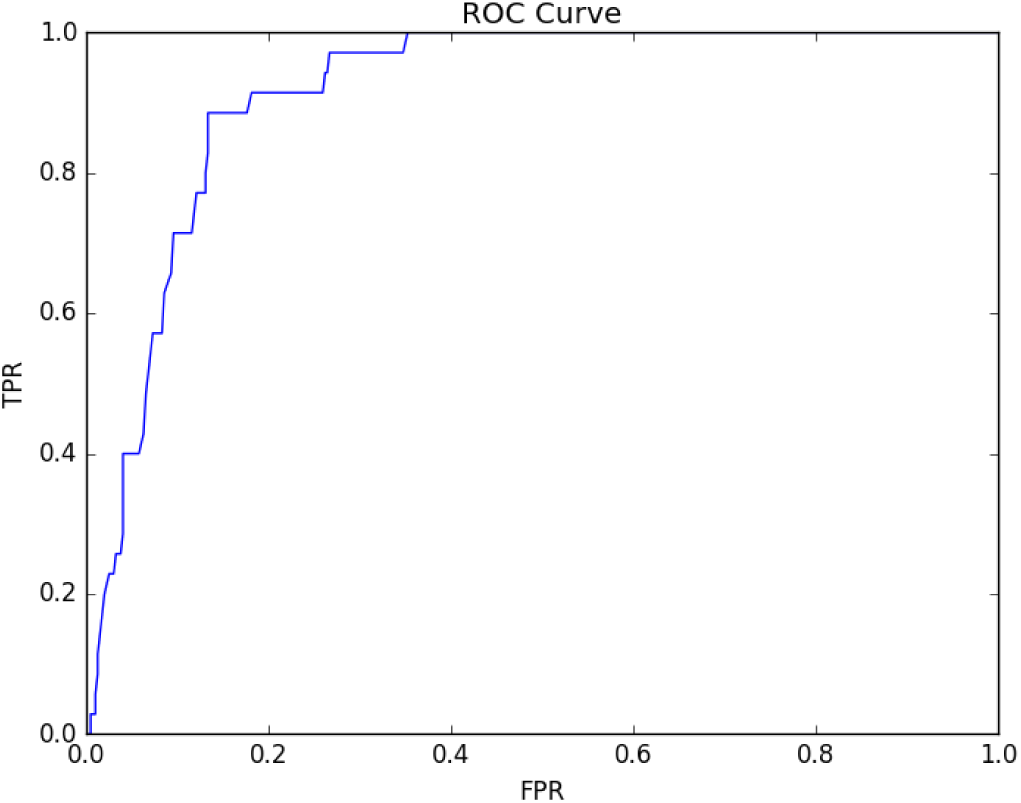
ROC curve of the binary outbreak classifier of the minimum discrepancy model.

**Figure 18:**
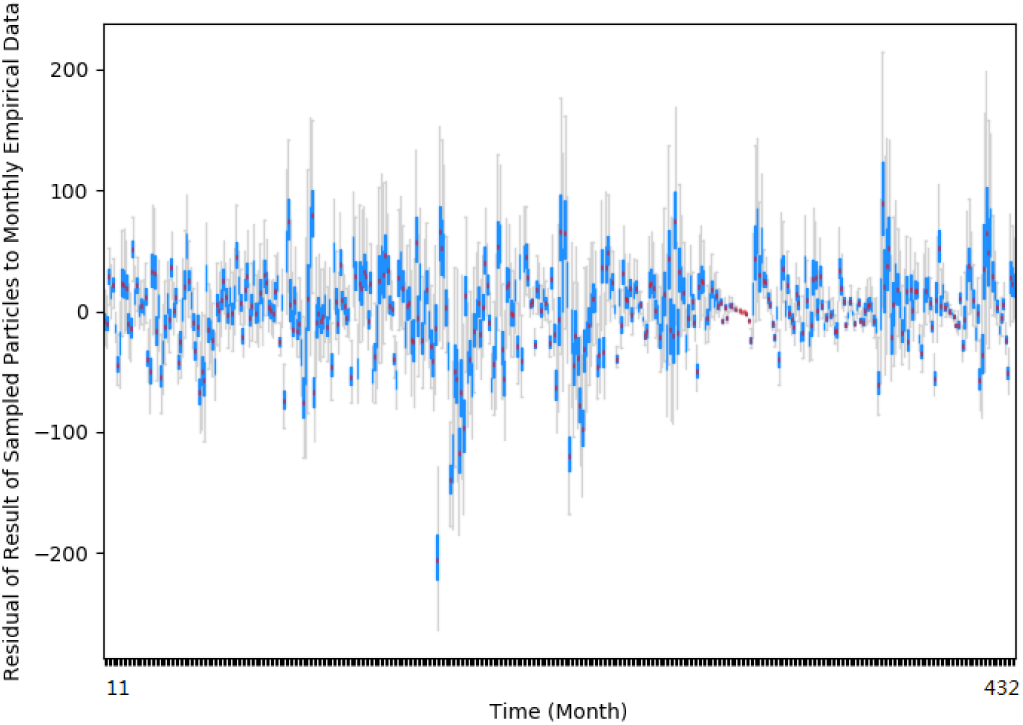
Boxplot of the residuals of results of prior prediction by sampled particles of the minimum discrepancy model.

### 3.4. Intervention with the minimum discrepancy model

The capacity of particle filtering to accurately estimate (sample from) the latent state of a pertussis model makes this technique capable of both estimating the entire latent state and using that estimation to project patterns of pertussis spread and waxing and waning of incidence in the near term, and to anticipate outbreak occurrence. The capacity to perform such state estimation within a mechanistic model also supports particle filtering models in more accurate simulation of the tradeoffs between intervention strategies, despite their counterfactual character.

In this section, we have implemented several experiments to simulate stylized public health intervention policies, based on the minimum discrepancy particle filtering pertussis model identified above. The stylized intervention strategies are characterized in an abstract way for demonstration purposes, and are typically performed before or at the very beginning of an outbreak. For simplicity, we examine them as a historical counterfactual that takes place at a certain historic context. Moreover, to support easy comparison with the baseline prediction results of the minimum discrepancy model absent any interventions, all of the intervention strategies are simulated starting at the start month of an outbreak (month 269) in this project. Moreover, in order to appropriately characterize how such techniques could be employed in public health scenario planning, we assume here that the start month of the intervention (month 269) is the “current time” in the scenario – that we wish to asses the effects of that intervention considering only the data available up to but not including month 269, and simulate the results of the intervention forward from that point. The baseline prediction result of the minimum discrepancy model absent any interventions is shown in Figure 13 (b). We examine below the impact of two stylized intervention policies – hygeine-enhancing and vaccination.

Figure 19 and Figure 20 display results from simulation of hygeine-enhancing intervention strategies [29] whose effects are characterized as decreasing the contact rate parameter by 20% and 50% when compared to its pre-intervention value, respectively. Similarly to the 2D histogram plot of the baseline prediction result shown in Figure 13 (b), the red dots represent the empirical data incorporated into the particle filtering model (here, up to just prior to the point of intervention); by contrast, the black dots represent empirical data not incorporated in the model, but presented for comparison purposes. It bears emphasis that because the interventions being characterized are counterfactual in character – i.e., did not in fact take place historically – the empirical data shown in black reflect the baseline context, which lacked an intervention of the sort simulated here. By comparing the hygeine-enhancing intervention results (see Figure 19 and Figure 20) with the baseline model result without intervention shown in Figure 13 (b) and the empirical data during the intervention period (the black-markers indicating historic data points lying after the triggering of the intervention, and not incorporated into the particle filtering model), we can see that, although the interventions are implemented in a stylized fashion, by virtue of the particle filter’s ability to estimate the underlying epidemiological state at the point of intervention through the transmission model, the particle filtered pertussis model is capable of using the estimated latent state to serve as the basis for probabilistically evaluating pertussis related intervention policies.

**Figure 19:**
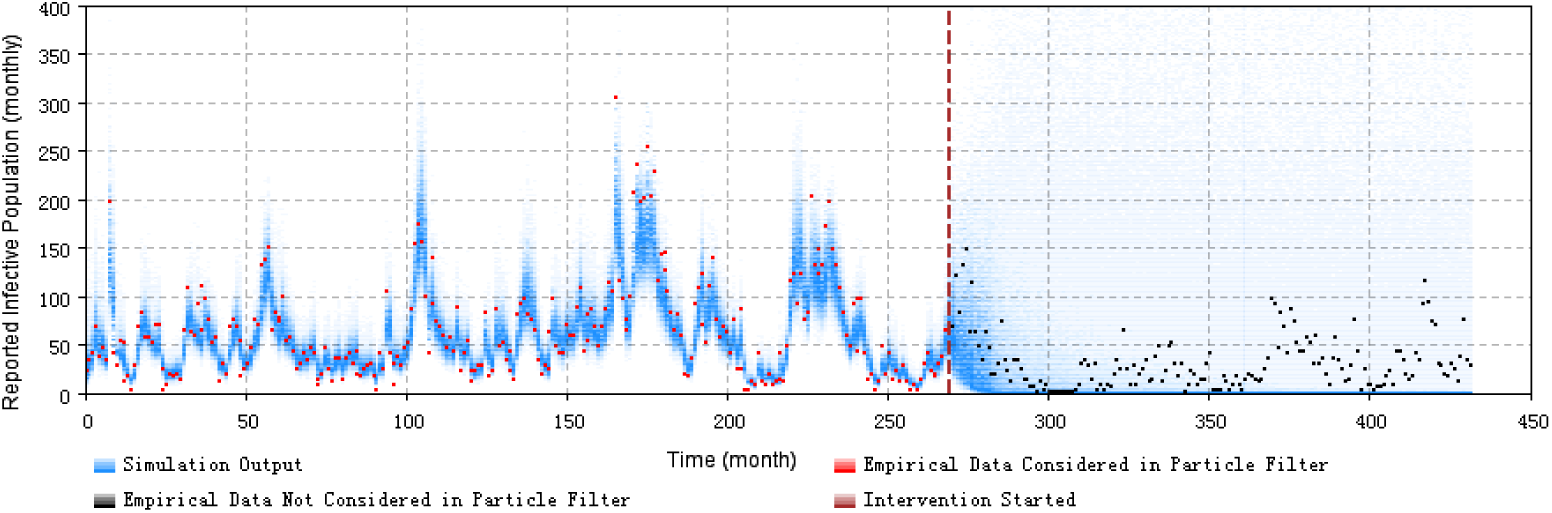
2D histogram of model-based projections of pertussis incident case counts when simulating a hygeine-enhancing intervention during a pertussis outbreak. This is realized by decreasing the contact rate by 20%.

**Figure 20:**
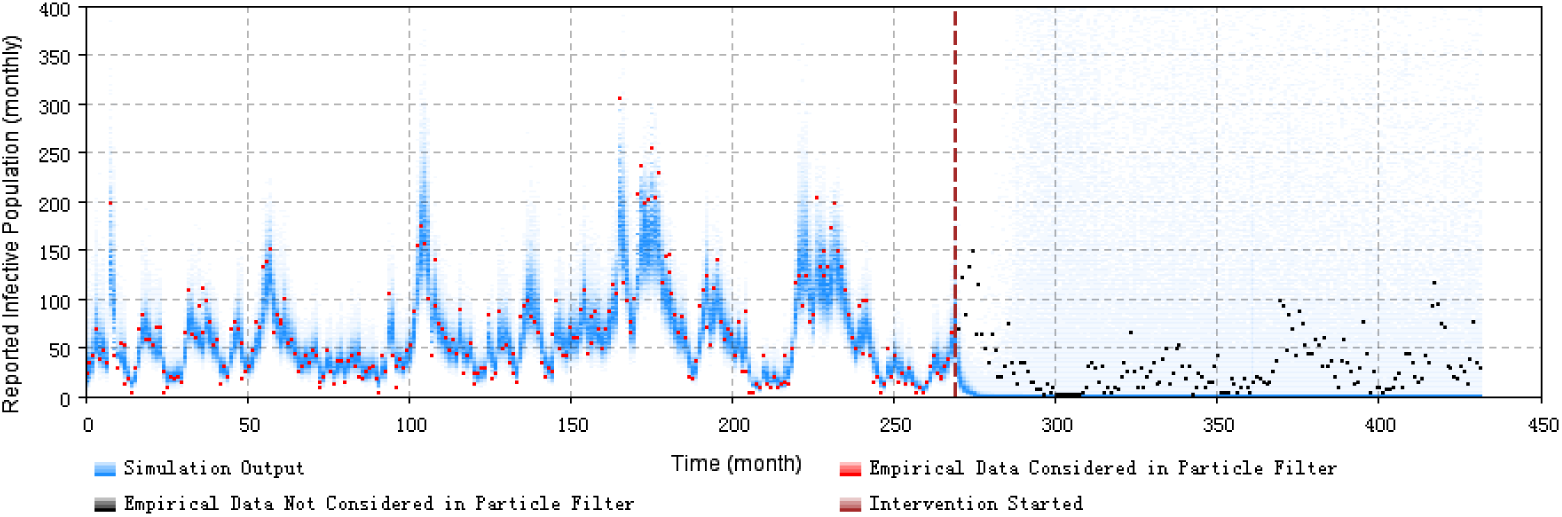
2D histogram of model-based projections of pertussis incident case counts when simulating a hygeine-enhancing intervention during a pertussis outbreak. This is realized by decreasing the contact rate by 50%.

To simulate an immunization intervention during a pertussis outbreak, a vaccination parameter is incorporated into the simulation model, so as to represent the fraction of the population whose immunity status is elevated as a result of the intervention. Specifically, recall that the pertussis model characterizes a chain of successively higher levels of vaccine-induced protection. This parameter specifies the fraction of the population that should be moved from their pre-vaccination classification – as characterized by the model compartment in which they reside – to the compartment representing the next higher level of vaccination (following vaccination). Figure 21 and Figure 22 show the results of the vaccination intervention. The layout and organization of the 2D histogram plots of the vaccination interventions with Figure 21 and Figure 22 mirrors that of the hygeine-enhancing plots of Figure 19 and Figure 20.

**Figure 21:**
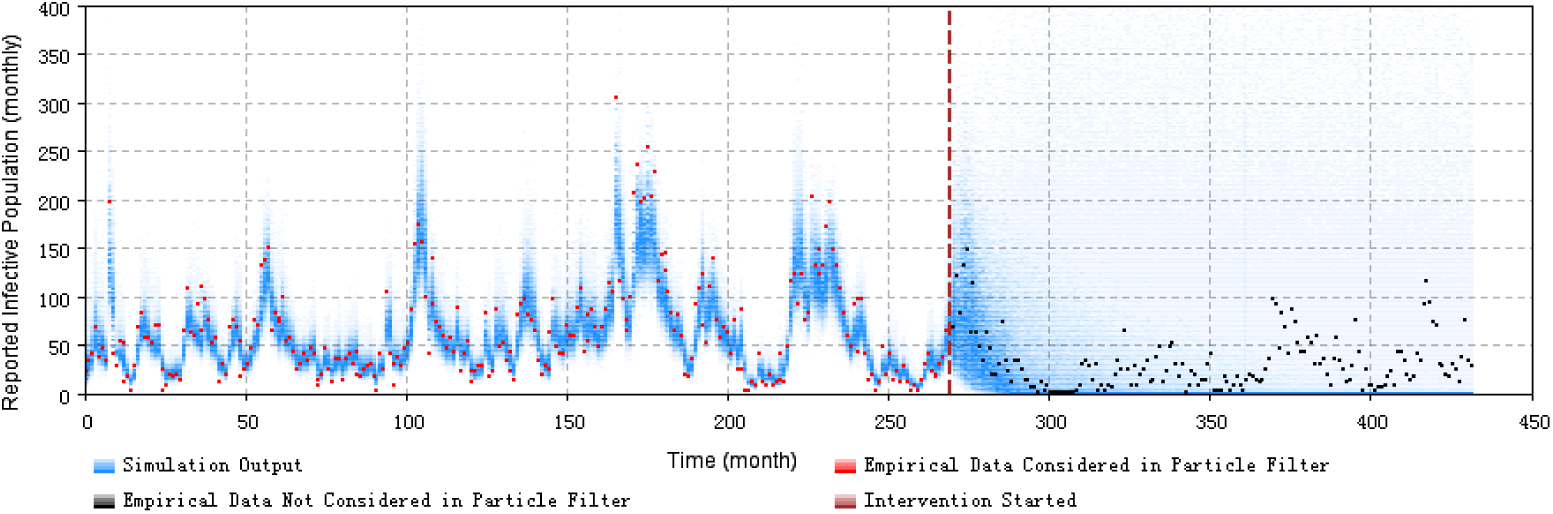
2D histogram of model-based projections of pertussis incident case counts when simulating an outbreak-response immunization campaign. This is realized by characterizing a stylized elevated vaccine-induced protection level among 20% of the population.

**Figure 22:**
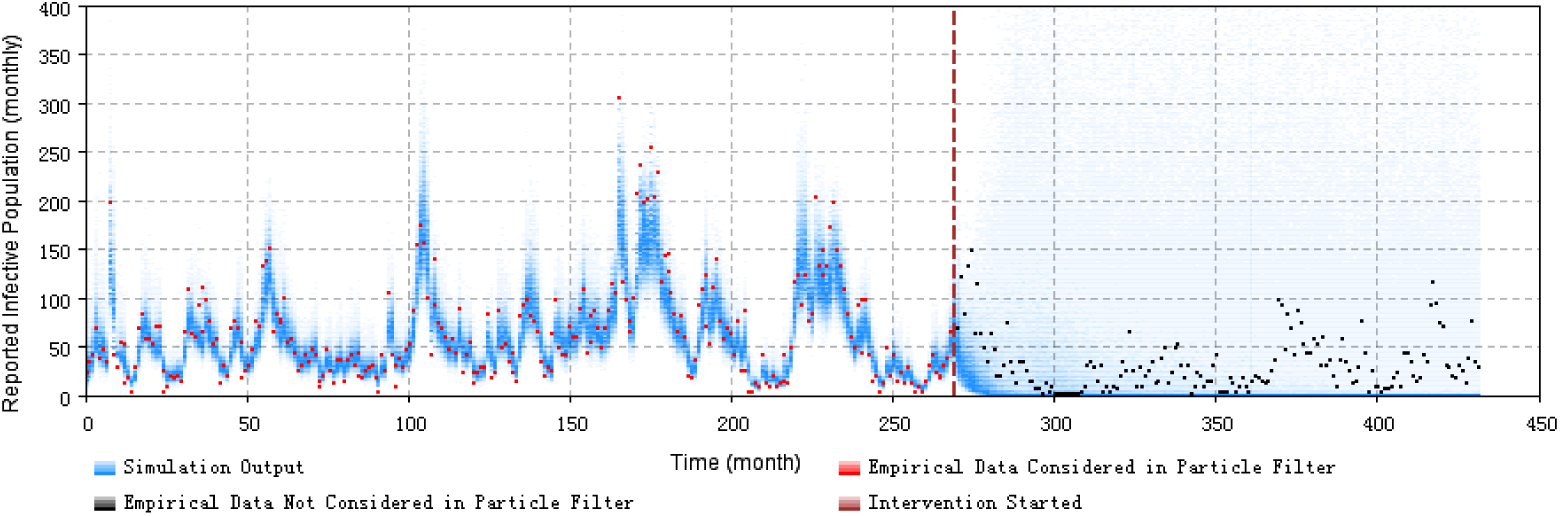
2D histogram of model-based projections of pertussis incident case counts when simulating an outbreak-response immunization campaign. This is realized by characterizing a stylized elevated vaccine-induced protection level among 50% of the population.

The results of pertussis interventions demonstrate that by virtue of its ability to estimate the underlying epidemiological state of the model (and thus the system characterized by that model), the use of particle filtering with pertussis models supports evaluation of public health intervention policies to prevent or control pertussis outbreaks.

## 4. Discussion and conclusion

This paper contributes a new method for anticipating, tracking, controlling and preventing pertussis outbreak patterns by integrating a particle filtering algorithm with a mechanistic pertussis compartmental model and empirical incidence data. This contribution represents the first time that particle filtering has applied to pertussis transmission dynamics, and demonstrates the great promise of this technique. The models examined here demonstrated a notable degree of accuracy in predicting pertussis dynamics over multi-month timeframes – the 2D histogram plots comparing the empirical data and samples from the posterior distributions of the particle filtering models’ projected monthly and yearly reported cases of pertussis indicates that the high probability density region of the model’s prediction of empirical data encompasses or lies near the historic data. The results of prediction analysis based on the minimum discrepancy model suggest that particle filtering approaches offer notable strengths in predicting of occurrence of pertussis outbreak in the subsequent month. Moreover, the discrepancy of the pertussis particle filtering model’s predictions vs. observed data is reduced by approximately 60% when compared with a traditional calibration model, demonstrating a significant enhancement in model prediction ability. Additionally, it bears emphasis that the calibrated deterministic model encounters marked difficulties in tracking the fluctuation of the outbreak pattern of the calibration model; by contrast, the particle filtered model is capable to tracking stochastic fluctuations associated with pertussis, while still mechanistically capturing the impacts of such stochastics on the latent underlying dynamics of susceptibles, exposed individuals, etc. Further to this point, it is of great significance to the success and promise of these methods that pertussis particle filtering models support effective estimation of the entire state of the pertussis transmission models – and thus the systems that they represent – during those periods when the empirical datasets are available, including latent states of strong interest, such as those associated with waning of natural immunity and differing levels of infection severity. Combined with the capability to perform outbreak projections, such particle filtering models can serve as powerful tools for understanding the current epidemiology of pertussis in the population, for projecting forward evolution of pertussis spread – including occurrence of outbreaks.

Beyond that, in a further contribution that also benefits strongly from the capacity to estimate latent state, this research further marks the first instance of research demonstrating the capacity to perform public health intervention experiments using particle filtered models.

Despite the strengths of these contributions, there remain a number of important limitations of this work, and priorities for future research. We briefly comment on several below.

This work investigated the performance of four particle filtering models, including an aggregate population model, a two age group-stratified population model, and 32 age group population models using – alternatively – a contact matrix derived from Hethcote (1997) [16] and (separately) a re-balanced contact matrix. Although the results of all four of these particle filtering models matched the empirical data quite well, the minimum discrepancy model proved to be the 2 age group age-stratified particle filtering model in which individuals in the child age group represent children in the first 5 years of life, and which incorporates both monthly and yearly empirical datasets. In this regard, it is notable that according to the mathematical deduction of the age structured population model introduced in [8] – and adapted to pertussis in this research – the model can simulate the aging rate (*c*_*i*_) more accurately with more age groups considered in the age-structured model. However, in this paper, the 32 age group particle filtering models fail to demonstrate improved performance – as measured by the discrepancy of model predictions from the empirical data – when compared with the two age group particle filtered models. We provide below some comments on possible reasons. Firstly, the stochastic processes considered in both the 32 age group age-structured model and the two age-group stratified model are different, especially in their characterization of the stochastic evolution of the contact rate. Secondly, the likelihood functions employed in this project – which are captured as the product of negative binomial density functions across all empirical datasets and sharing a common dispersion parameter – may be too naïve to capture the difference between the age groups within the empirical datasets. Thirdly – and perhaps most significantly – as the number of age groups increase, the the state space dimensionality of the particle filtering models increases dramatically. This latter issue must be considered in light of the limitations of the particle filtering algorithm, particularly the fact that the particle filtering method employing the condensation algorithm may encounter problems in high-dimensional systems. In such systems, the probability density functions would be more involved; addressing this using the functional form of the likelihood functions employed may require high dispersion, due to the difficulty of representing the details of the multivariate likelihood function using the product of simple probability density functions. Research is needed into more effective multivariate likelihood function design. The relationship between the the nominal state space dimensionality and the number of particles required for effective particle filtering also merits additional research, particularly in light of observed limitations in the benefit of particle filtering for high dimensional models [13]. Finally, when comparing the discrepancy for distinct models, our lack of normalization for the count of datasets used may lead to artificially stacking the comparison against the 32 age group model; while the 32 age group does not exhibit markedly better discrepancy against the monthly aggregate observations than does the 2 age group model, this consideration suggests that it may be stronger than the yearly discrepancy numbers would suggest.

It is also worth emphasizing the critical role that stochastic process noise within the state space models plays within successful particle filtering, and the practical challenges associated with managing such noise. The stochastics associated incorporated into the model represent a composite of two factors. Firstly, there is expected to be both stochastic variability in the measles infection processes (e.g., those that are prominent for small incident case counts) and some evolution in the underlying transmission dynamics in terms of changes in mixing and the reporting rate. Secondly, incorporation of such stochastic variability into the particle filtered model allows for characterization of uncertainty associated with respect to model dynamics – reflecting the fact that both the observations and the model dynamics share a high degree of fallibility, and allowing a requisite variety in the distribution over particle states, such that the particle filtered model is more open to correction by new observations. While results in both the estimation and prediction periods are sensitive to the degree of stochastics involved, such model stochastics impact the particle filtered model in distinct distinct ways during these periods. Taking into account these influences, the investigations demonstrated the importance of keeping the noise in the particle filtering models controlled within a proper range, by tuning the parameters of diffusion coefficients in the stochastic processes related to the Brownian motion. The need to characterize and tune stochastic noise effectively can impose limits on the speed with which particle filtering models can be prepared for a new sphere of application.

The initial values of the age-structured population models in this paper are estimated both manually and by the particle filtering algorithm. Specifically, the population distribution among the different age groups are tuned manually, while the population distribution among different compartments within a given age group is estimated by the particle filtering algorithm by setting the initial values of compartments in a proper range following a uniform distribution, but maintaining a total number of individuals for that age group across the compartments. Especially in building the 32-age-groups particle filtering models, much time and efforts is dedicated to estimation of the population distribution among the latent states.

While application of particle filtering to pertussis dynamics is not without its challenges, the approach examined here demonstrates great promise for creating models that are automatically kept abreast of the latest evidence, for understanding the underlying epidemiology of pertussis in the population – including the balance of the population at varying levels of immunity – for projecting forward pertussis dynamics and outbreak prediction over a year’s time, and for evaluation of counter-factual interventions. The results of this paper – which represents both the first application of particle filtering to pertussis, the first to demonstrate the capacity to accurately predict pertussis outbreaks in the pre-vaccination era, and the first to use particle filtering to assess the tradeoff between public health applications – suggest that particle filtering may represent an important element in the arsenal of public health tools to address the increasingly difficult challenge of controlling pertussis in the context of vaccine hesitancy and waning of both natural- and vaccine induced-immunity.

## 5. Future work

The growing risk of pertussis outbreaks triggered by combinations of vaccine hesitancy and waning dynamics from earlier generations of pertussis vaccination, has elevated the urgency and prominence of questions about the rate at which immunity to pertussis wanes, especially about vaccine effectiveness over time [30, 31, 32, 33]. We identify here two notable needs for future work responsive to such dynamics Firstly, there is a keen need for application of the models presented here to data and dynamics from the vaccination era. While vaccination elements of the models discussed here are only glancingly tapped by this research (in the context of demonstrating capacity to reason about the effects of a stylized immunization intervention), because of their incorporation into the existing model structure, extension of this work to vaccine-era dynamics should require only limited changes to the models involved.

A second need relates to the fact that we choose to employ a constant value for the waning of immunity that is drawn from [16]. In the future, to arrive at more informed parameter estimates for models and to contribute to discussion concerning the empirical rate at which vaccine-induced as well as (separately) naturally induced pertussis immunity wanes, and drawing on the success of our past work in this area [34], we propose to formally estimate the value of waning immunity for the the particle filtered pertussis models from a posterior distribution within Particle Markov Chain Monte Carlo (PMCMC) techniques, by incorporating empirical data on reported pertussis cases from the vaccination era.

These and other lines of future work offer substantial promise in extending the already strong potential demonstrated here for using mechanistic transmission models informed by the machine learning approach of particle filtering to contribute to enhancements in pertussis prevention and control by providing a tool to improve understanding of underlying complex epidemiology of pertussis, to anticipate pertussis dynamics in the population, and to rigorously assess the tradeoffs between counterfactual intervention tradeoffs in light of uncertainties in both model and empirical data.

## Appendix A. The introduction of mathematical models

In this compartmental model of pertussis in the pre-vaccination era, the total population is divided into 8 distinct epidemiological classes. Newborns enter directly into class *S* of fully susceptible individuals. If a fully susceptible individual contacts an infective individual and is successfully transmitted pertussis, this previous susceptible person becomes infectious and enters the class (state) *I* of full infectives. Infective individuals in state *I* of have full cases of pertussis, with all of the usual symptoms. When individuals recover from the state *I* of infectives, they achieve full immunity and enter state *R*_4_. In this state, they are fully protected and can not be infected by pertussis. However, as time goes by, their immunity wanes and they enter into a less strong immunity class of *R*_3_. When individuals in class *R*_3_ are exposed to an infective, they are assumed to return to the highest immunity class of *R*_4_ without becoming infectious. Otherwise, their immunity keeps fading, and they enter to the relatively lower immunity class of *R*_2_. When a person in the class of *R*_2_ is sufficiently (re-)exposed to an infective for transmission to occur, the infected individual enters the *I*_*w*_ state with weak infectivity. Individuals in the *I*_*w*_ class have the weakest infective capability to infect a susceptible. After they recover, the individuals in class of *I*_*w*_ then secure the highest immunity, (re-) entering the class of *R*_4_ from which they originally waned. By contrast, if people in the class of *R*_2_ are not re-exposed to the infectives, their immunity continues waning, and they enter the minimally immune class of *R*_1_. Similarly, if a person in class *R*_1_ is re-exposed to an infective, this person gets infected with mild infectivity and enters the class of *I*_*m*_. Individuals in the class of *I*_*m*_ have a higher infectious capability compared with those in the class of the weak infective (*I*_*w*_), but exhibit a lower infectious capability compared to the fully infective individuals in *I*. When recovered, the individuals in class *I*_*m*_ enter the class *R*_4_ again. If the individuals in the class of *R*_1_ are not re-exposed, they eventually lose all of their immunity and move back to the class of *S* whence they originated at birth. Given the presence of multiple infection states (*I, I*_*w*_, *I*_*m*_) as well as multiple levels of immunity (*R*_1_, *R*_2_, *R*_3_, *R*_4_), three invariants bears noting. Firstly, regardless of the pre-existing level of immunity, following recovery from an infection, that individual always returns to the full level of natural immunity (*R*_4_). Secondly, in any of the recovered states (*R*_1_, *R*_2_, *R*_3_, *R*_4_), immunity continues to wane absent re-exposure. Thirdly, as the level of immunity is reduced, the severity of resulting infectiousness rises, with no infectiousness being possible at all from exposure in states *R*_3_ and *R*_4_.

## Appendix B. Proof of the n-square grows of the unknown parameters

In this part, we prove that the unknown parameters grows with n-squared with the total number of age groups in the model. The contact matrix has been introduced, which is

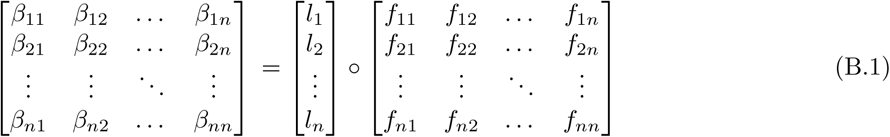

where ^∘^ indicates the Hadamard (element-wise) product; the parameter of *l*_*i*_ (1 *≤ i ≤ n*) is the contact rate of age group *i*. In this research, the *l*_*i*_ is known variables; the parameters of *f*_*ij*_ (1 ≤ *i* ≤ *n*, 1 ≤ *j* ≤ *n*) indicates the fraction of the age group *j* of the contact rate of the age group *i*.

The *f*_*ij*_ are normally unknown. And the total number of *f*_*ij*_ is *n*^2^. However, there are two relationships under this method. One relationship is that the sum of the fraction to all the age groups of the age group (e.g. *i*) is 1.0. The other relationship, related to the characteristics of balance of the contact matrix, is that the total contacts of the age group *i* to the age group *j* should be equal to the total contacts of the age group *j* to the age group *i*. Based on these two relationships, two equations could be generated as follows:

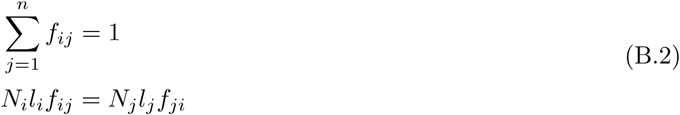

the total number of equations in Equation (B.2) is 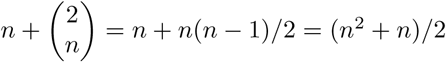. Finally, in this method of calculating the contact matrix, the number of unknown parameters is (*n*^2^ − *n*)/2. It indicates that the number of the unknown parameters grows in n-squared with the total number of age groups (n) in the model.

## Appendix C. The mathematical deduction of the force of infection with re-balanced contact matrix

In the beginning, we introduce the method of calculating the basic contact matrix which is balanced already and with one unknown parameters. Before introduced, we import a mixing parameter, denoted as *ϵ*. The mixing parameter *ϵ* determines where mixing occurs on a scale from fully associative – persons only contact with the individuals in the same age group (e.g. *ϵ* = 0) and random mixing – the contact among the total population is homogeneous (e.g. *ϵ* = 1.0). Then, the fraction of the average persons that an individual in age group *i* that contact with the persons in the age group of *j*, which is the parameters of *f*_*ij*_ in the contact matrix are represented as follows:

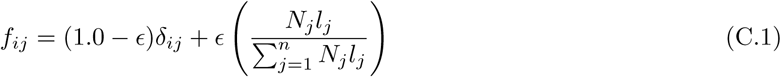

where *δ*_*ij*_ is the identity matrix. And the elements in the contact matrix is *l*_*i*_*f*_*ij*_.

The total contacts of age group *i* to age group *j* (*N*_*i*_*l*_*i*_*f*_*ij*_) equal the total contacts of age group *j* to age group *i* (*N*_*j*_*l*_*j*_*f*_*ji*_), in this basic contact matrix. And the only unknown parameter is *ϵ*. However, in general, the mixing parameter related to each age group should be different. For example, the mixing parameter of *ϵ* of young children in school age maybe lower than the *ϵ* of the little baby, because the children in the school age contacts more to their peers in the school than the other groups, while the little baby contacts more with their parents or care-taker than the other babies. Thus, in the next step, we expend the mixing parameter *ϵ* to a vector, where each element represents the mixing parameter of each related age group *ϵ*_*i*_.

Then, in this method of calculating the contact matrix with a vector of mixing parameters, the equation of *f*_*ij*_ is listed as follows:

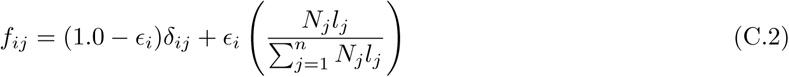

Similarly, the elements in this contact matrix with a vector of mixing parameters are *l*_*i*_*f*_*ij*_. It is notable that the total contacts between any two age groups calculated based on this contact matrix are unbalanced. Specifically, the number of total contacts of age group *i* to age group *j* is 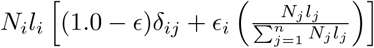, while the number of the total contact of age group *j* to age group *i* is 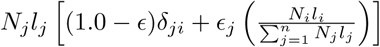. In general, the mixing parameters of any two age groups are not the same. Thus, the total numbers of contacts calculated by this contact matrix between any two age groups are not always the same.

To make the contact matrix balanced, we have employed the method introduced in [21] to re-balance the contact matrix. A parameter, denoted as Δ_*ij*_, is imported to represent the ratio of the number of total contacts between any two different age groups (*i* ≠ *j*) (for the same age group, the total number of contacts are always the same). Then, the equation of Δ_*ij*_ is:

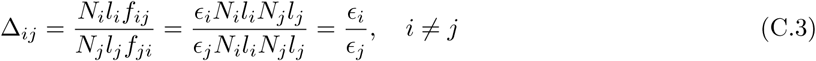

Then, the main idea of re-balancing the contact matrix is to extend the vector of contact rates (the elements of the contact rates are denoted as *l*_*i*_) to a new matrix of contact rates *l*_*ij*_. The elements in the matrix of contact rates *l*_*ij*_ represent the number of persons in the age groups *j* that a person in the age group *i* could contact in average. Then, according to [21], the equations of *l*_*ij*_ and *l*_*ji*_ could be defined separately:

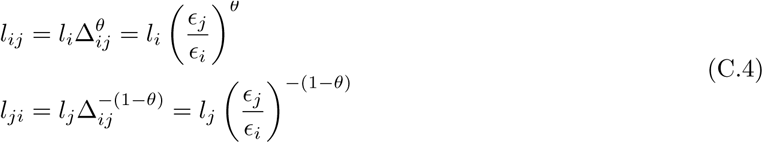

where *θ* is the re-balanced parameter.

Because both *l*_*ij*_ and *l*_*ji*_ represent the same matrix, a relationship could be generated, which is *l*_*ij*_ = *l*_*ji*_. Then, we could get the value of the parameter of *θ* (*θ* = 0.5). Substitute the value of *θ* (*θ* = 0.5) to Equation (C.4), the matrix of contact rate – *l*_*ij*_ could be generated as follows:

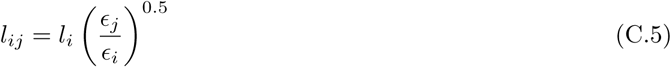

Finally, the element of contact matrix *l*_*ij*_*f*_*ij*_ and force of infection *λ*_*i*_ are:

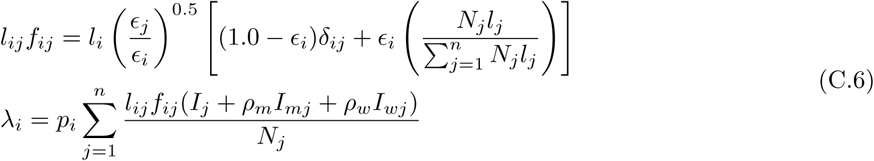

where *p*_*i*_ is the transmission probability of age group *i*.

## Appendix D. The state space models in Particle filter implementation

The mathematical system dynamics models are employed as the governing equations underlying the state space model. Then each particle at time *k*, noted as 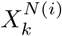, represents a complete copy of the system states at that point of time. Except for the basic states in the mathematical model – pure Ordinary Differential Equations (ODEs), models of infection transmission are often related to more complex dynamics – such as parameters evolving according to stochastic processes.

In this paper, we employ the identity method introduced in the previous contribution of [8] to let some constant parameters in the pure mathematical models change dynamically. Specifically, if the parameter varies over the entire range of positive real numbers, we treat the natural logarithm of this parameter as undergoing a random walk according to a Wiener Process (Brownian Motion) [25, 26, 8]. Otherwise, if the parameter varies over the range [0,1], we characterize the logit of this parameter as also undergoing Brownian Motion.

### Appendix D.1. The aggregate model (n = 1)

In the aggregate model, the individuals contact with the infectious (in the stocks of *I*_*w*_, *I*_*m*_ and *I*) homogeneously. Then, three stochastic processes are considered in the implementation of the aggregate particle filtering model. The first is the transmissible contact rate linking infectious and susceptible persons, which is represented by the parameter *β*. The second is also with respect to the disease reporting process. Specifically, a parameter – representing the probability that a given pertussis infectious case is reported *C*_*r*_, and a state *I*_*k*_ – calculating the accumulative pertussis infectious cases per unit time (per Month in this project) – are implemented. The final part is the Poisson process associated with the incidence of infection. This process reflects the small number of cases that occur over each small unit of time – Δ*t* (0.01 in this model). We also treat the natural logarithm of the transmissible contact rate (denoted by *β*) and the logit of *C*_*r*_ as undergoing a random walk according to a Wiener Process (Brownian Motion) [25, 26, 8]. It is notable that we assume the individuals under the medium infectious (*I*_*m*_) and weak infectious (*I*_*w*_) also have the probability to be confirmed and reported. The rates of the medium infectious (*I*_*m*_) and weak infectious (*I*_*w*_) that have symptoms are also considered as *ρ*_*m*_ and *ρ*_*w*_. Finally, the state space model of the aggregate pertussis particle filtering model is listed as follows:

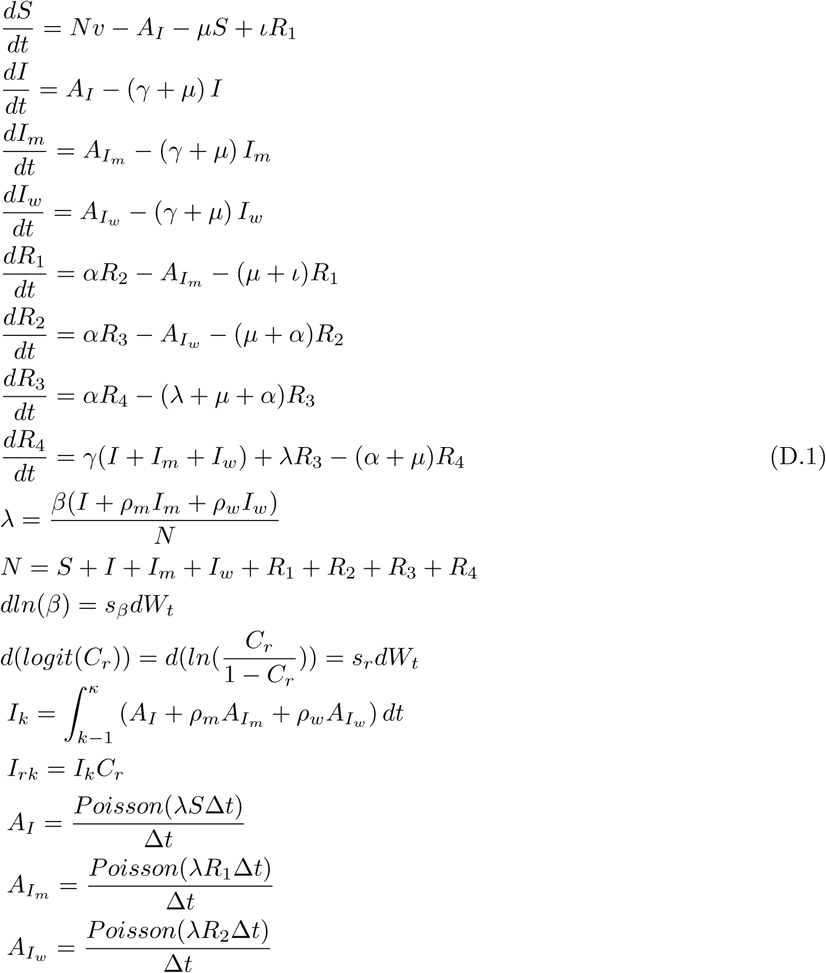

The parameters related to the transmission of pertussis in this model are referred from the research of Hethcote (1997) [16]. The demographic parameters of this model are got from the Annual Report of the Saskatchewan Department of Public Health [20] and the age pyramid of Saskatchewan [19]. Then, the parameters of the pertussis aggregate state space model – Equations (D.1) are specified in Table D.2, while the initial values of the stocks are listed in Table D.3.

**Table D.2:** Table showing the value of parameters in the pertussis aggregate particle filtering model.

| Parameter | Description | Value | Units |
| --- | --- | --- | --- |
| $\gamma^{-1}$ | mean time for infectives to recover from pertussis | 21 | Day |
| $v$ | birth rate of the total population | 0.03 | 1/Year |
| $\mu$ | death rate of the total population | 0.03 | 1/Year |
| $N$ | total population | 863,545 | Person |
| $\iota^{-1}$ | mean time to lose immunity from the stock of $R_1$ to $S$ | 10 | Year |
| $\alpha^{-1}$ | mean time to lose immunity from $R_i$ down to $R_{i-1}$ | 5 | Year |
| $\rho_m$ | the relative infectivities of the individuals in the stock $I_m$ | 0.5 | Dimensionless |
| $\rho_w$ | the relative infectivities of the individuals in the stock $I_w$ | 0.25 | Dimensionless |
| $s_\beta$ | the diffusion parameter of $\ln(\beta)$ | 0.5 | Dimensionless |
| $s_r$ | the diffusion parameter of $\ln(\frac{C_r}{1-C_r})$ | 0.05 | Dimensionless |

**Table D.3:** Table showing initial values of the stocks in the pertussis aggregate particle filtering model.

| Parameter | Value | Unit |
| --- | --- | --- |
| $S_0$ | Uniform[5000, 30000) | Person |
| $I_0$ | Uniform[500, 5000) | Person |
| $I_{m0}$ | 1000 | Person |
| $I_{w0}$ | 2500 | Person |
| $R_{10}$ | Uniform[10, 10000) | Person |
| $R_{20}$ | 10000 | Person |
| $R_{30}$ | 20000 | Person |
| $R_{40}$ | $N - S_0 - I_0 - I_{m0} - I_{w0} - R_{10} - R_{20} - R_{30}$ | Person |
| $\beta$ | Uniform[5,100) | Person/Month |
| $C_r$ | Uniform[0,0.2) | Dimensionless |

### Appendix D.2. The age-structured model of 2 age groups (n = 2)

The mathematical model with two age groups is employed as the base model of the state space model of the age-structured model with 2 age groups. Then, the pure ODEs model – mathematical model – is extended by several stochastic processes. Except for the similar three stochastic processes considered in the aggregate state space model – the infectious contact rate of the child age group (denoted as *β*_*c*_), the report rate of pertussis cases (denoted as *C*_*r*_), and the Poisson process related to the incidence of the infectious – two other stochastic processed are also considered. These two stochastic processes are related to the parameter of the multiplier of the adult age group model (*M*_*a*_) of the infectious contact rate and the fraction of children’s infectious contacts that occur with other children (*f*_*cc*_). Specifically, the natural logarithm of the multiplier of the infectious contact rate of the adult age group (*M*_*a*_) and the logit of *f*_*cc*_ are treated as undergoing a random walk according to a Wiener Process (Brownian Motion) [25, 26, 8]. Finally, the state space model of the pertussis age-structured model of 2 age groups is listed as follows:

**Table D.4:** Table showing the value of parameters (only related to the demographic model and stochastic processes) in pertussis two-age-groups particle filtering model.

| Parameter | Description | Value | Units |
| --- | --- | --- | --- |
| $v_a$ | birth rate of the adult age group | 0.034 | 1/Year |
| $N_c$ | the population of the child age group | 98743 | Person |
| $N_a$ | the population of the adult age group | 764802 | Person |
| $\omega$ | the aging rate from child to adult age group | 0.2 | Dimensionless |
| $s_\beta$ | the diffusion parameter of $\ln(\beta)$ | 0.5 | Dimensionless |
| $s_r$ | the diffusion parameter of $\ln(\frac{C_r}{1-C_r})$ | 0.05 | Dimensionless |
| $s_{M_a}$ | the diffusion parameter of $\ln(M_a)$ | 0.2 | Dimensionless |
| $s_{cc}$ | the diffusion parameter of $\ln(\frac{f_{cc}}{1-f_{cc}})$ | 0.15 | Dimensionless |

**Table D.5:**
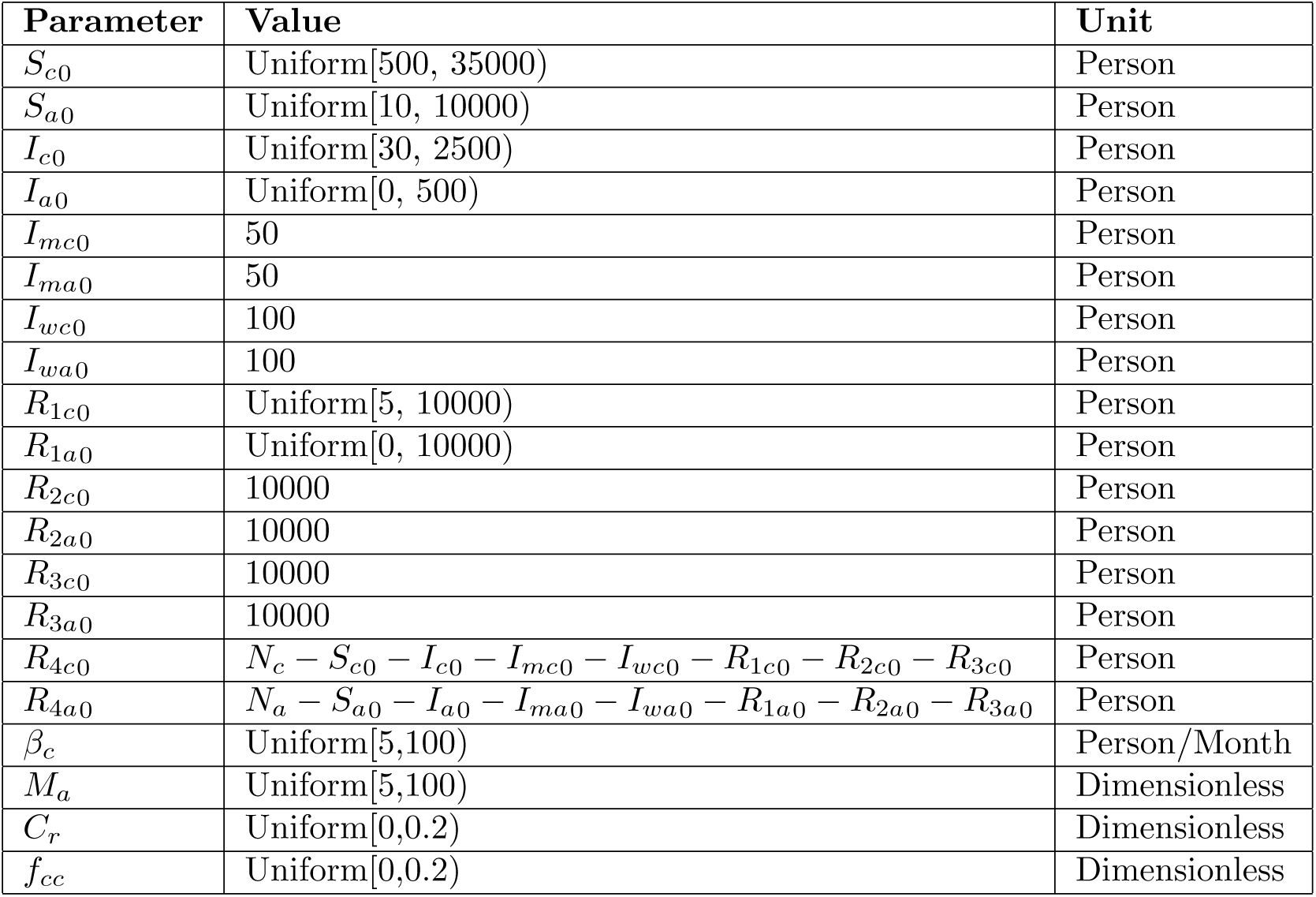
Table showing initial values of the stocks in the pertussis two-age-groups particle filtering model.

| Parameter | Value | Unit |
| --- | --- | --- |
| $S_{c0}$ | Uniform[500, 35000) | Person |
| $S_{a0}$ | Uniform[10, 10000) | Person |
| $I_{c0}$ | Uniform[30, 2500) | Person |
| $I_{a0}$ | Uniform[0, 500) | Person |
| $I_{mc0}$ | 50 | Person |
| $I_{ma0}$ | 50 | Person |
| $I_{wc0}$ | 100 | Person |
| $I_{wa0}$ | 100 | Person |
| $R_{1c0}$ | Uniform[5, 10000) | Person |
| $R_{1a0}$ | Uniform[0, 10000) | Person |
| $R_{2c0}$ | 10000 | Person |
| $R_{2a0}$ | 10000 | Person |
| $R_{3c0}$ | 10000 | Person |
| $R_{3a0}$ | 10000 | Person |
| $R_{4c0}$ | $N_c - S_{c0} - I_{c0} - I_{mc0} - I_{wc0} - R_{1c0} - R_{2c0} - R_{3c0}$ | Person |
| $R_{4a0}$ | $N_a - S_{a0} - I_{a0} - I_{ma0} - I_{wa0} - R_{1a0} - R_{2a0} - R_{3a0}$ | Person |
| $\beta_c$ | Uniform[5,100) | Person/Month |
| $M_a$ | Uniform[5,100) | Dimensionless |
| $C_r$ | Uniform[0,0.2) | Dimensionless |
| $f_{cc}$ | Uniform[0,0.2) | Dimensionless |

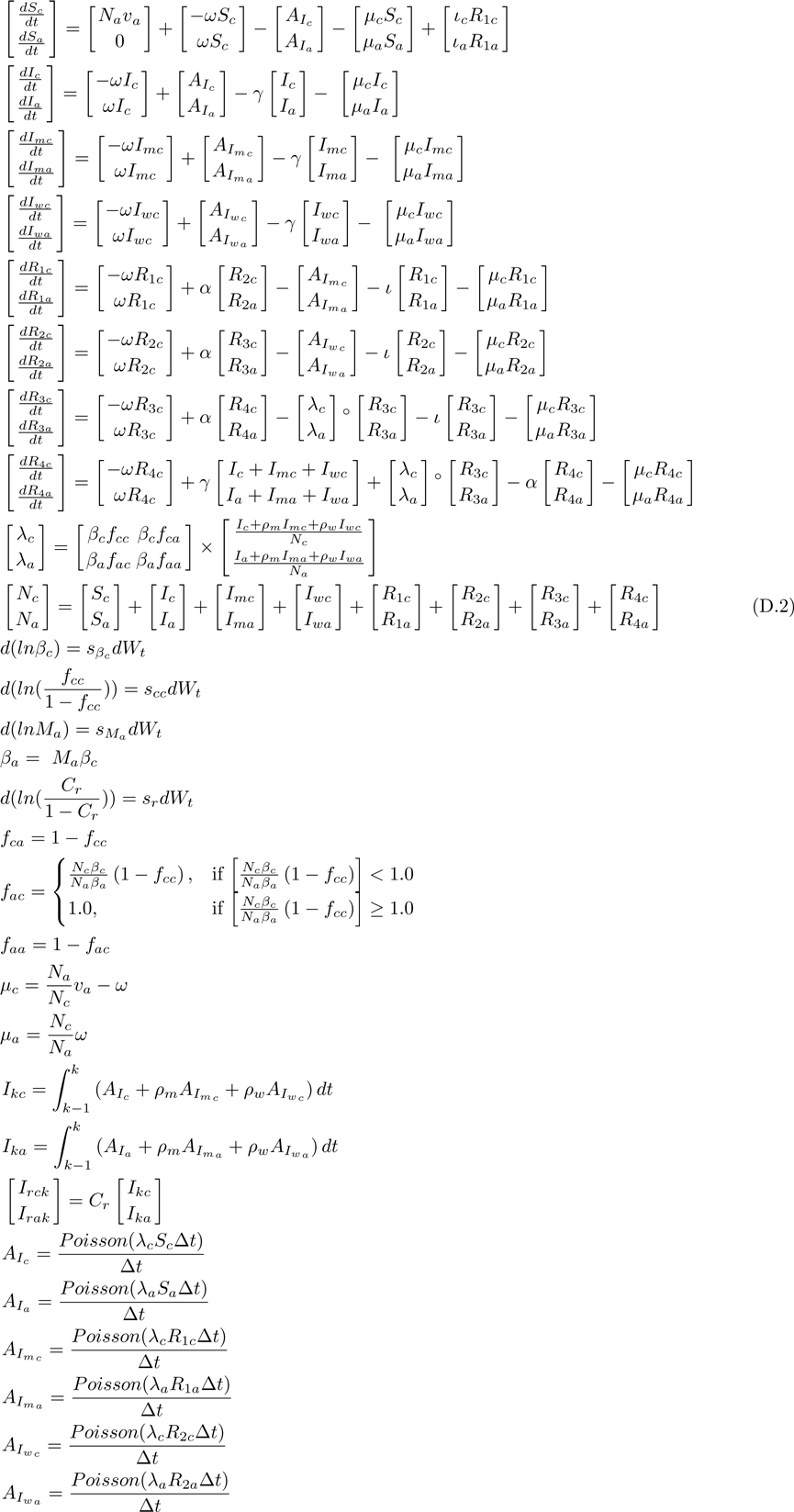

In this paper, we have built a two-age-group particle filtering model, where the individuals in the age group of “child” are from newborn up to the end of 4 years. The parameters with constant values related to the pure compartmental model (*γ, ι, α, ρ*_*m*_ and *ρ*_*w*_) in the two-age-group pertussis model are the same as the aggregate model. All these parameters and the parameters related to the demographic model and the stochastic processes of the two-age-group particle filtering model are listed in Table D.4. The initial values of each stocks in this two-age-group particle filtering model are listed in Table D.5.

### Appendix D.3. The age-structured model of 32 age groups (n = 32) with the Hethcote contact matrix

We employ the pure ODEs model – the age-structured model of 32 age groups introduced in the paper of Hethcote (1997) [16] as the base model. Similarly, three stochastic processes are added to the base model as the state space model. These three stochastic processes are related to the Poisson process related to the incidence of infectious, the contact rate of the first age group and the reporting process of the pertussis cases. Similarly, the natural logarithm of the parameter related to contact rate of the first age group (denoted as 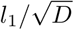) and the logit of the report rate (denoted as *C*_*r*_) are treated as undergoing a random walk according to a Wiener Process (Brownian Motion) [25, 26, 8]. :

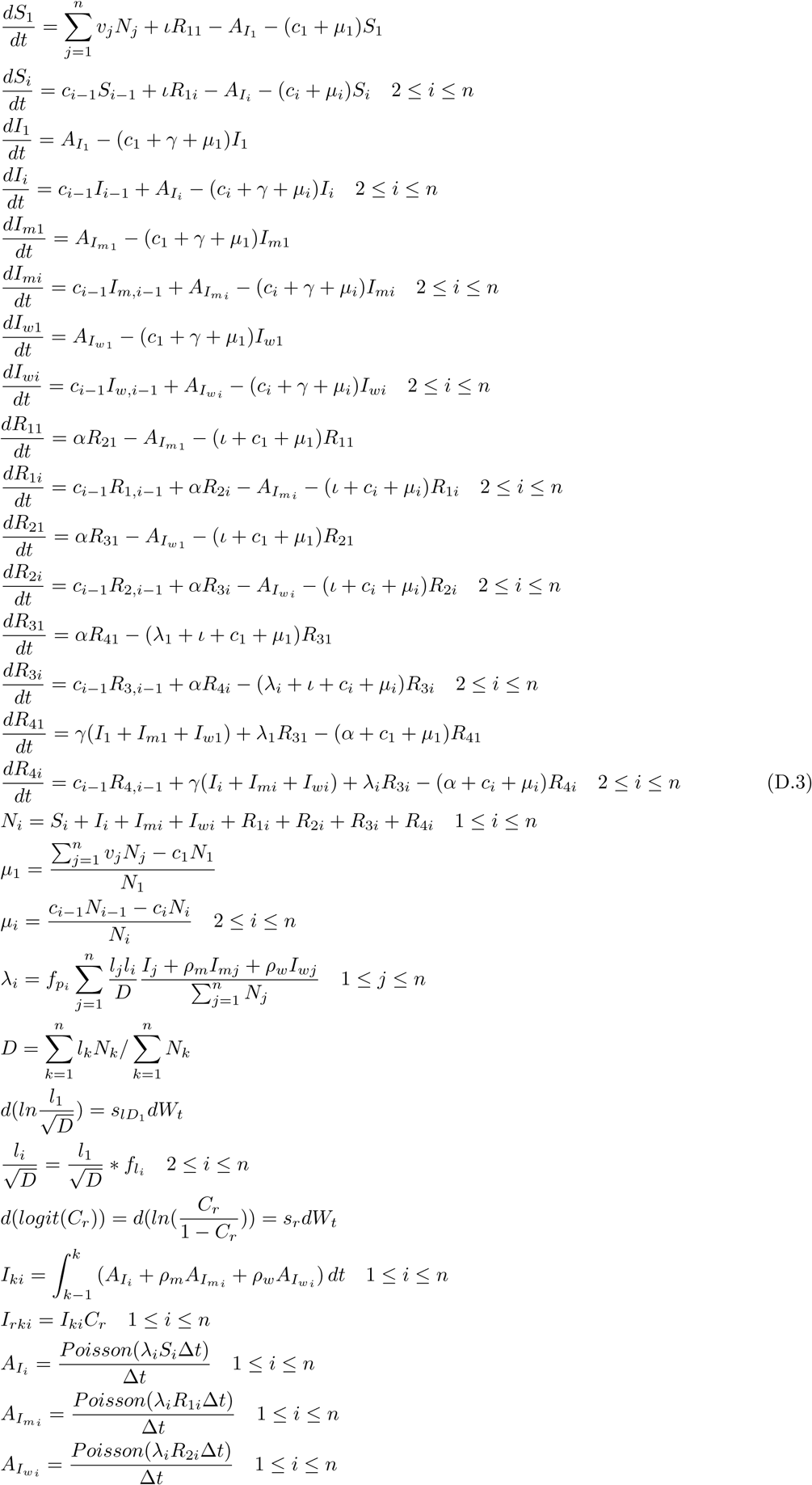

The values of the parameters are the same as the ones listed in the aggregate particle filtering models and two-age-group particle filtering model, and the initial values of the stocks in this particle filtering model are listed in the Table D.6.

### Appendix D.4. The age-structured model of 32 age groups (n = 32) with re-balanced contact matrix

The age-structured model of 32 age groups with re-balanced contact matrix are employed as the base model of the state space model of the age-structured particle filtering model of 32 age groups with re-balanced contact matrix. The mathematical equations of state space model are listed as follows:

**Table D.6:**
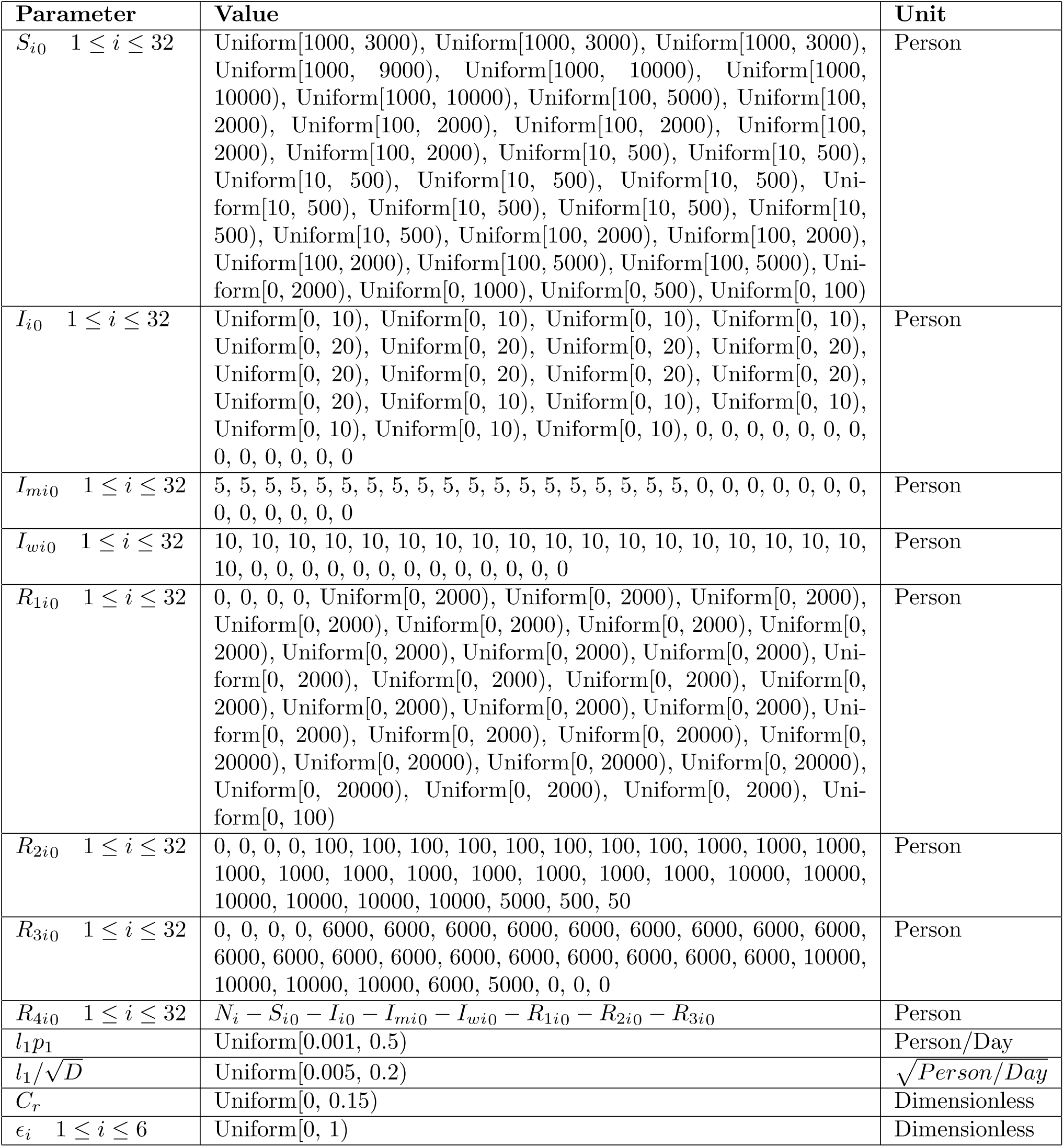
Table showing initial values of the stocks in the pertussis 32-age-groups particle filtering models.

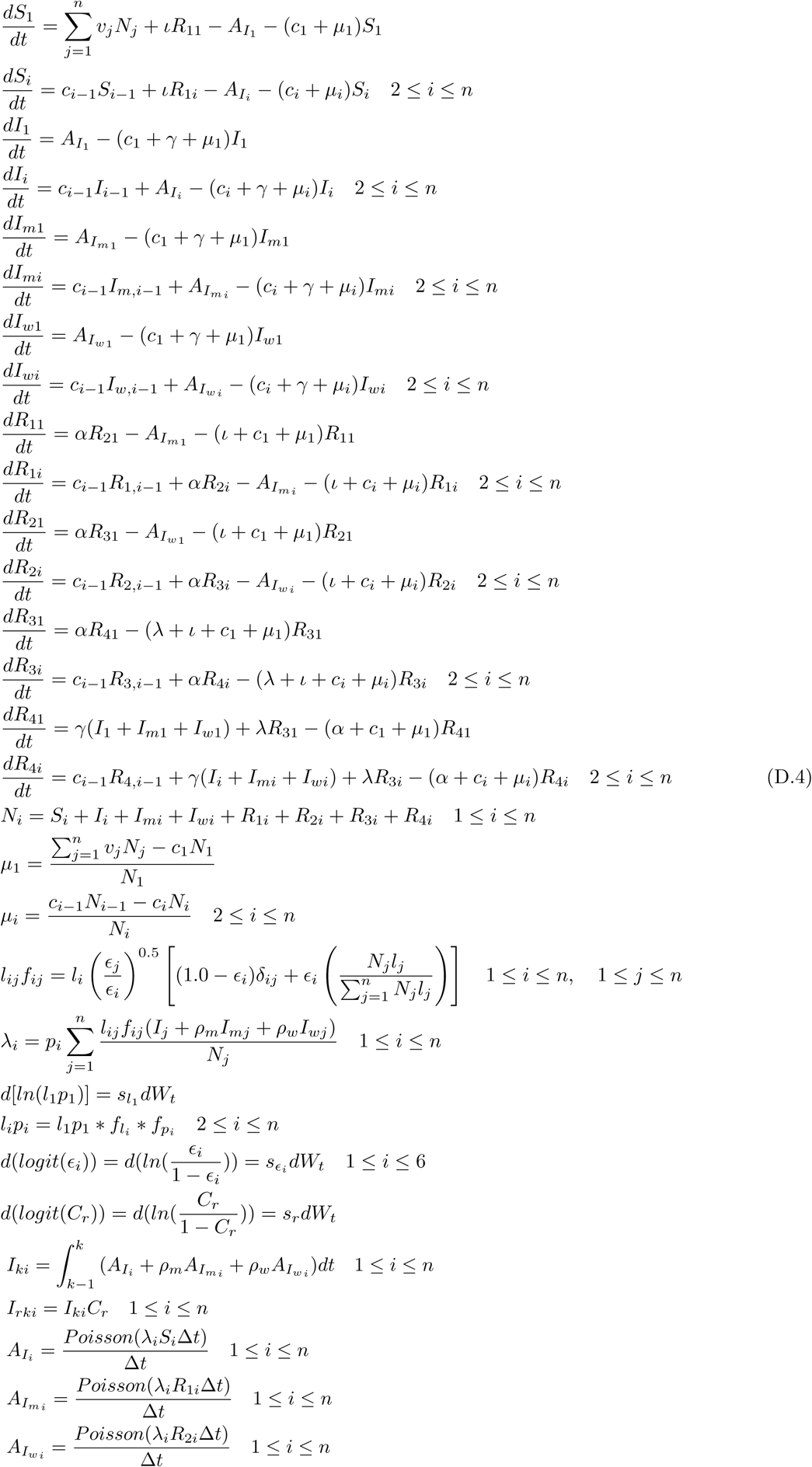

The values of the parameters are the same as the ones listed in the aggregate particle filtering models and two-age-group particle filtering model, and the initial values of the stocks in this particle filtering model are listed in Table D.6.

## Appendix E. Further introduction of split the pertussis yearly reported cases to each age group

The yearly empirical data related to multiple age categories are available from year 1925 to 1956 [20]. During the process in preparing the yearly empirical data for the two-age-group and 32-age-group particle filtering models (the yearly empirical data divided into 6 groups), we need to split the data in some age categories in the original datasets [20] due to two reasons. The first reason is because the division of the age group in empirical dataset does not match the division of the age groups in the pertussis particle filtering models. Specifically, from year 1926 to 1941, we need to split the reported pertussis cases in age category “1-6 years” in age 5 proportionally (four fifths goes to the “1-4 years” age group, and one fifth goes to “5-9 years” age group); from year 1942 to 1955, we need to split the reported pertussis cases in age category “5-14 years” in age 10 proportionally (half goes to the “5-9 years” age group, and half goes to “10-14 years” age group). The second reason is because there is a category in the empirical yearly dataset of “age not stated”. Thus, we need to split the counts in this category to corresponding age groups in the particle filtering models proportionally (based on the proportion calculated by the age categories has labeled age clearly).

